# The evolution of human SIRT1 for multi-specificity is enabled by active-site adaptation

**DOI:** 10.1101/2020.01.12.903153

**Authors:** Adi Hendler, Eyal Akiva, Mahakaran Sandhu, Dana Goldberg, Eyal Arbely, Colin J. Jackson, Amir Aharoni

## Abstract

Many enzymes that catalyze protein post**-**translational modifications (PTM) can specifically modify multiple target proteins. However, little is known regarding the molecular basis and evolution of multi-specificity in these enzymes. Here, we used bioinformatics and experimental approaches to investigate the molecular basis and evolution of multi-specificity in the sirtuin-1 (SIRT1) deacetylase. Guided by bioinformatics analysis of SIRT1 orthologues and substrates, we identified and examined key mutations that have occurred during the evolution of human SIRT1. We found that while these mutants maintain high activity toward conserved histone substrates they exhibit a dramatic loss of activity toward acetylated p53 protein that appeared only in Metazoa. These results demonstrate that active site substitutions in SIRT1 are essential to enable the inclusion of p53 substrate. Our results are consistent with a model in which promiscuous ancestral activity can provide an evolutionary starting point from which multispecificity toward a number of cellular substrates can develop.

## Introduction

The evolution of protein-protein interaction (PPI) networks from simple mono-cellular eukaryotes (e.g. yeast) to a multicellular organism (e.g. human) is often characterized by a dramatic increase in the number of cellular proteins and PPI complexity (Evlampiev and Isambert, 2008; Jancura et al., 2012; Jin et al., 2013; Liang et al., 2014). Conserved hub proteins, located at the heart of these PPI networks, must maintain or expand their multi-specificity to allow the recognition of a growing number of partners. Currently, little is known regarding the molecular basis for multi-specificity in PPI networks and how hub proteins (Han et al., 2004) evolve to recognize a large and diverse set of protein partners. In some PPIs, hub-partner interactions occur through a defined consensus recognition sequence found in different partners. Network expansion in these cases can take place by the appearance of a recognition sequence or structural motif in nascent partners during evolution (Cino et al., 2013; Kim et al., 2009; Moldovan et al., 2007). However, in many PPI networks, shared recognition sequence/motifs are not found in partners and the molecular basis for hub-partner interactions is unknown. In these PPIs our understanding of the molecular basis and evolution of hub-partner interactions is very limited.

Many enzymes that catalyze the post-translational modification (PTM) of proteins are located at the heart of complex PPI networks that expand through natural evolution (Beltrao et al., 2013). These enzymes catalyze the specific attachment or removal of different functional groups including phosphate, acetyl, methyl and ubiquitin. A substantial number of PTM-catalyzing enzymes found in human cells exhibit remarkable ability to recognize many protein substrates and catalyze the modification of specific target residues. Such specificity allows the regulation of a variety of essential cellular processes including DNA replication, transcription or signal transduction (Beltrao et al., 2013; Deribe et al., 2010).

One of these enzymes is the human SIRT1 (hSIRT1) that belongs to the large family of sirtuin enzymes (Finkel et al., 2009). The sirtuins are NAD+-dependent deacetylases (Blander and Guarente, 2004; Haigis and Guarente, 2006) and the founding member of this family, yeast Sir2 (ySir2), was initially identified in *S. cerevisiae* (Greiss and Gartner, 2009; Imai et al., 2000). These enzymes are conserved from bacteria to humans and their overexpression in several eukaryotes was shown to increase organism’s life span (Cohen et al., 2004; Howitz et al., 2003; Kanfi et al., 2012). The hSIRT1 is the mammalian ortholog of ySir2 and is the best studied of the human sirtuins (Lavu et al., 2008). The number of SIRT1 substrates expanded dramatically during evolution from several acetylated lysine positions on histones for ySir2 to hundreds of acetylated sites in different substrates of hSIRT1 (Chen et al., 2012; Gil et al., 2017; Rauh et al., 2013). The involvement of hSIRT1 in central biological processes like DNA repair, stress resistance, apoptosis and aging, occurs through the deacetylation of central regulatory substrates including p53, histones, FOXO, HSF1 and NFkB leading to modulation of their biological activities (Daitoku et al., 2004; Westerheide et al., 2009; Yeung et al., 2004). Despite extensive efforts, little is known regarding how hSIRT1 recognizes the target acetyl-lysine in a variety of different substrates and no consensus sequence in hSIRT1 substrates was identified (Blander et al., 2005; Garske and Denu, 2006; Rauh et al., 2013). Moreover, it is unknown how multi-specificity is enabled in hSIRT1 and whether specific substitutions took place during SIRT1 evolution enabling the expansion of its substrate repertoire. Thus, SIRT1 provides an excellent model system for the study of the molecular basis and evolution of multi-specificity in enzymes with a diverse set of substrates and cellular functions.

Here, we developed and applied a bioinformatics-experimental workflow to identify substitutions in hSIRT1 enabling its high degree of multi-specificity. Our bioinformatics approach exploits natural variation in hSIRT1 homologs, ancestral sequence reconstruction and the expansion of SIRT1 substrate repertoire to identify changes in key residues that may enable hSIRT1 multi-specificity. Guided by this bioinformatics analysis, we generated different hSIRT1 mutants containing substitutions that are located at the vicinity of hSIRT1’s active site. Examination of these mutants revealed high activity with the highly conserved acetylated histones but a dramatic reduction in activity toward p53, which only appeared after divergence of fungi and animalia, relative to the WT protein. Our data suggest that active site substitutions that occurred during natural SIRT1 evolution are essential for the high degree of hSIRT1 multi-specificity.

## Results

### A bioinformatics workflow for identifying candidate positions important for SIRT1 multi-specificity

To identify a limited number of candidate positions that may be important for hSIRT1 multi-specificity, we combined sequence similarity networks (SSN), which are graphical summaries of sequence similarities shared between homologous proteins (Akiva et al., 2017; Atkinson et al., 2009), multiple sequence alignments (MSA), phylogenetic trees and phylogenetic profiles focusing on the deacetylase (DAC) domain of SIRT1. The construction of a reliable MSA of SIRT1 homologs requires homogenous sampling of SIRT1 sequence space. Thus, we first delineated the sequence-similarity boundary that differentiates SIRT1 family members from the large sirtuin superfamily. We constructed an SSN of the entire sirtuin superfamily including 9521 sequences and then mapped specific sirtuins, documented in the literature, to identify the different sirtuin classes (SIRT1 to 7). This representation enabled the delineation of SIRT1 subgroup containing 1107 sequences that can be clearly separated from other sirtuin groups (**Fig. S1**). Next, we constructed an MSA of the SIRT1 family, including 103 unique sequences, by sampling it using criteria like homogenous taxonomic sampling, fully sequenced genomes and manual inspection (**Fig. S2**).

To follow the evolution of substrate recognition in SIRT1 by identifying semi-conserved amino-acids, we constructed a Bayesian inference phylogenetic tree based on the SIRT1 multiple sequence alignment (**Fig. S3**). Using the combination of the protein tree, taxonomy and the alignment, we identified positions that vary in their conservation levels – from universally conserved positions to ones that are conserved only in specific branches (partially conserved residues) and may be associated with SIRT1 adaptation to multi-specificity (**Fig. 1** and **Fig. S2**). Overall, this analysis allowed us to focus on 8 different positions (**Fig. 1**). Interestingly, four of these positions are located at the vicinity of hSIRT1’s active site, two positions are located at the Rossman fold domain and two positions at the zinc-binding domain (**Fig. 2**).

**Fig. 1:**
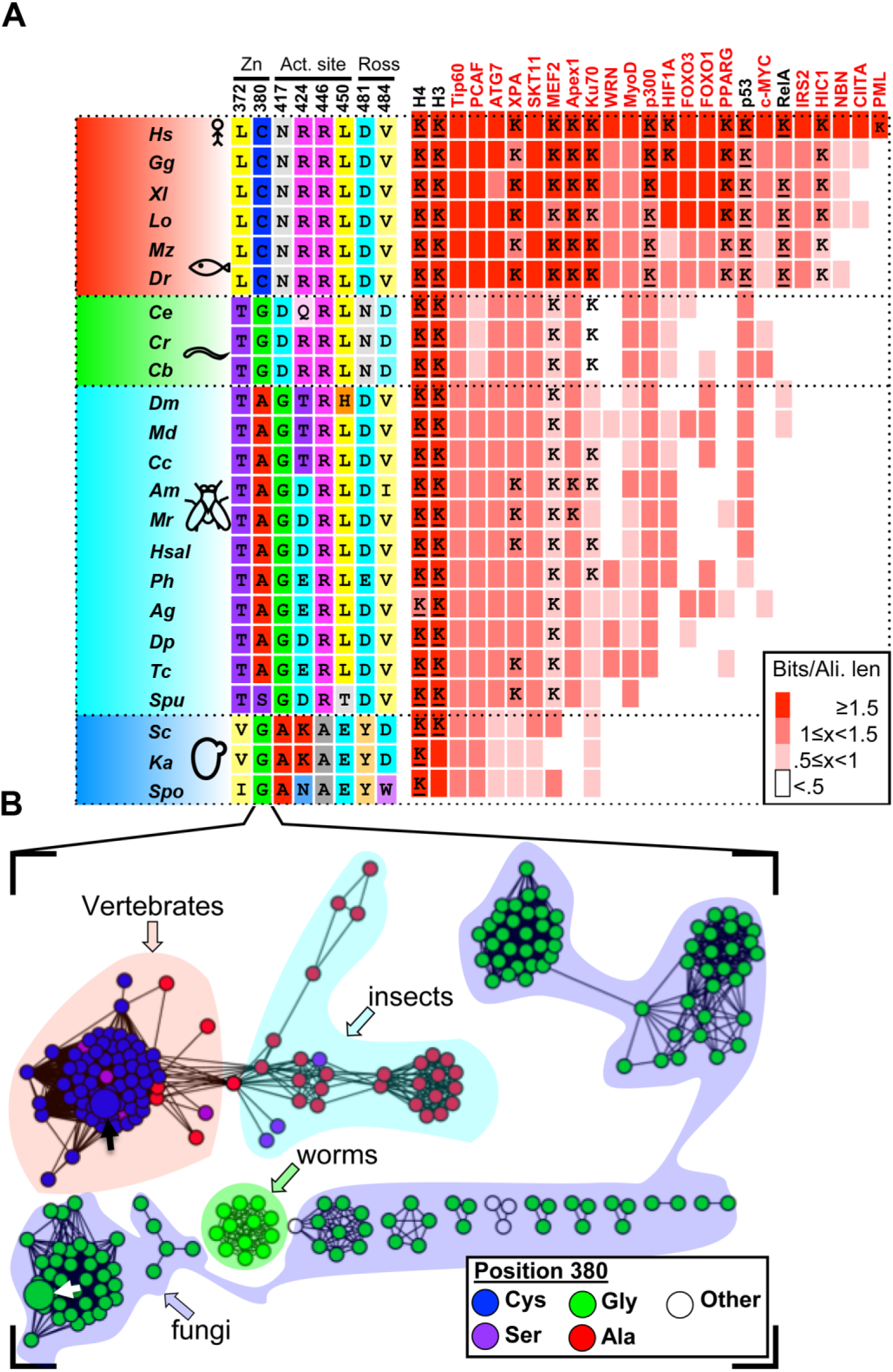
SIRT1 evolution is correlated with substrate repertoire expansion. (**A**) A phylogenetic tree of SIRT1 orthologs is shown on the left, representing major evolutionary branches of eukaryotic taxa: vertebrates (red), invertebrates I (green, mainly mullusca and nematodes), invertebrates II (cyan, mainly Ecdysozoa [insects]) and fungi (blue). A tree with organism names and branching probabilities is shown in **Fig. S3**. The positions selected for substitutions (see text and **Figs. 2-3**) demonstrate conservation that is associated with the major evolutionary transitions. Right, a matrix depicting the conservation of 25 hSIRT1 substrates in eukaryotes. The substrates tested in this study are shown in black headings. The conservation level of the orthologs, measured by Bit score divided by alignment length is color-coded from white (low conservation) to red (high conservation). The appearance of orthologs in the different organisms is cross-validated by the inParanoid (Sonnhammer and Östlund, 2015) and EggNOG (Huerta-Cepas et al., 2016) databases, as well as literature review and manual examination. In cases where the acetylated lysine is known for the hSIRT-1 human substrate, a “K” is given in the relevant matrix cell. Underlined “K” represent the substrate column tested in this study. (**B**) A sequence similarity network of 103 likely orthologs of hSIRT-1. Each node represents proteins that share 90% sequence identity and above. The BLAST *E*-value cutoff for edge inclusion is 1*10^−80^. The organic layout in Cytoscape (Shannon et al., 2003) is used to layout the network. Node colors represent the residues that correspond with hSIRT1’s position 380, and the taxonomy is shown as semi-transparent shapes, color-coded as in the tree in panel A.

**Figure 2:**
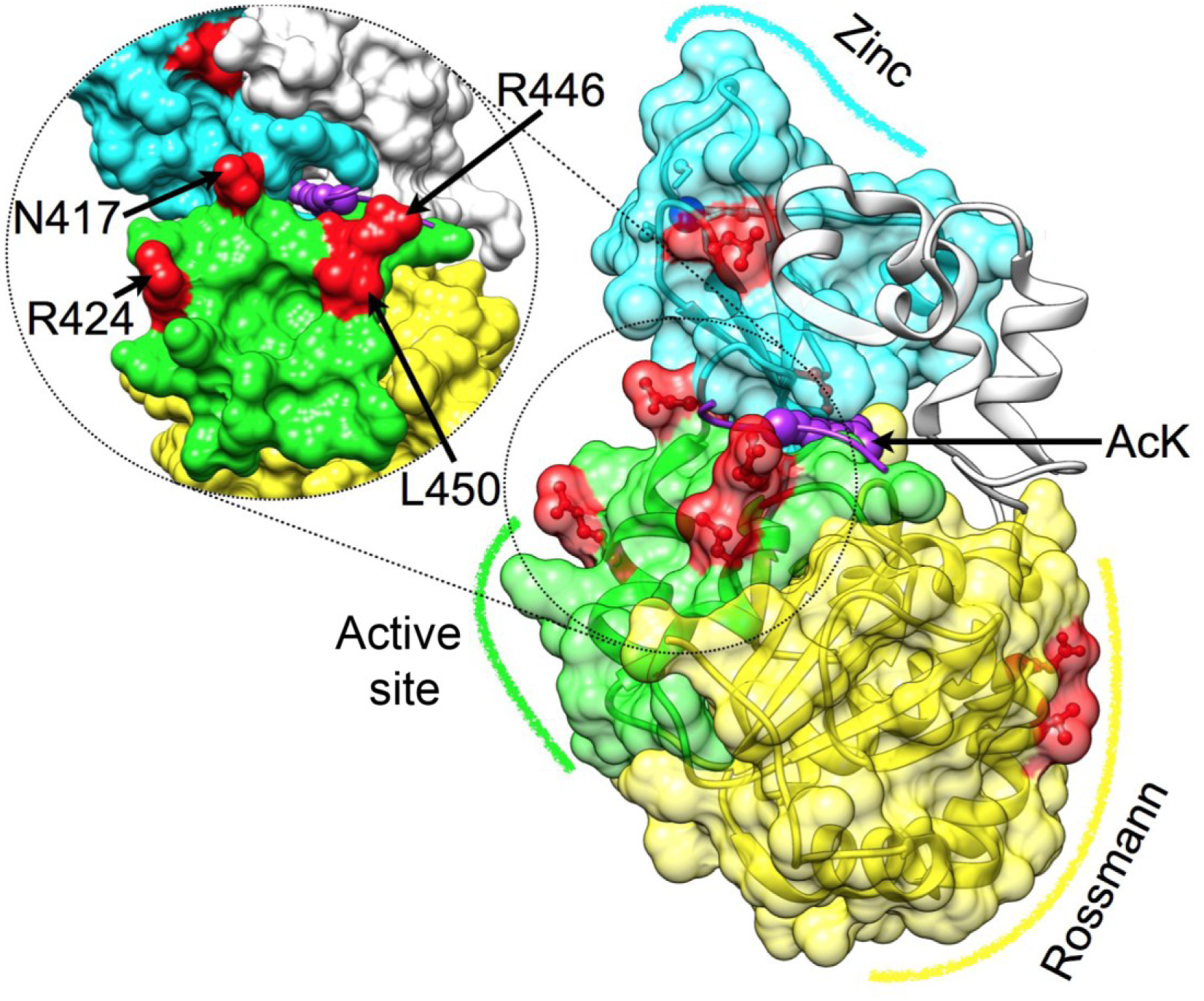
Representation of the DAC of hSIRT1. Three major structural regions in hSIRT-1 are shown (PDB 4KXQ), along with the specific positions that were substituted (red) Rossmann (yellow, positions 481 and 484), the active site region (green, positions 417, 424, 446 and 450, also tilted and enlarged on the left) and the Zinc binding domain (cyan, positions 372 and 380, the zinc ion is in blue). The peptide with the acetylated lysine is shown in purple, its approximate position is based on Sir2-p53 complex (PDB code 2H4F) (Davenport et al., 2014). This position highlights the proximity between positions located at the active site region and substrate specificity.

### Evolutionary dynamics of SIRT1 substrate expansion

The identification of evolutionary branches in which the orthologs of hSIRT1 substrates have emerged can be used for correlating between natural substitutions in SIRT1 and substrate expansion (**Fig. 1**). To this end, we first collected 25 experimentally validated substrates of hSIRT1 for which their acetylated lysine is partially known (Chen et al., 2012; Rauh et al., 2013) (**Table S1**). Next, we performed phylogenetic profiling for these substrates, *i.e.* assessing the presence or absence of orthologs for all 103 organisms from which SIRT1 alignment was constructed (**Fig. S2**). To assign a confidence score for their presence or absence, we combined data from several established methods (e.g. EggNOG (Huerta-Cepas et al., 2016), InParanoid(Sonnhammer and Östlund, 2015)), as well as manual inspection and literature search (see Methods). In addition, we generated MSA of these orthologs, aimed at evaluating the conservation level of the target acetyl-lysine in SIRT1 substrates (for examples see **Fig. S4**). As expected, we found that histones are completely conserved SIRT1 substrates while p53 and RelA are examples of substrates that appeared later in evolution in complex eukaryotes associated with SIRT1 substrate expansion (**Fig. 1**). In almost all cases, we identified the appearance of the substrate, followed by fixation of a lysine residue in a location that agrees with the acetyl lysine in the human substrate (**Fig. 1**). This classification of modern and primitive substrates, as well as lysine conservation, allowed us to select specific substrates for our experimental analysis (see below) and correlate between the substitutions identified in hSIRT1 and substrate analysis (**Fig. 1**).

### Experimental analysis of hSIRT1 mutants indicates intramolecular coevolution

Based on the bioinformatics and structural analysis described above, we generated, expressed and purified 13 hSIRT1 mutants (V1-V13) containing 1 to 4 substitutions in different regions of the DAC domain (**Figs. 2-3**). The deacetylation activity of the purified mutants was measured with generic acetylated lysine using the fluor de-lys (FDL) assay(Wegener et al., 2003) and compared to the WT hSIRT1. This analysis allowed us to reveal intramolecular coevolution between different hSIRT1 residues. Examination of V1-V3 mutants, containing mutations in the vicinity of hSIRT1 active site (**Fig. 2**), showed that the deleterious effect of L450E (V2) is compensated by R446A (V1) in the double mutant (V3) indicating significant coevolution between these two positions (**Fig. 3**). We found that the N417A and R424E mutations (V4) located at the vicinity of the active site does not dramatically affect hSIRT1 activity. However, the combination of N417A/R424E and R446A/L450E (V5) leads to a significant increase in SIRT1 deacetylase activity by ∼3 fold relative to the WT (**Figs. 3** and **S5**). Thus, V5 containing 4 active site vicinity mutations that mainly originates from fungi SIRT1 orthologues (**Fig. 1**) exhibits significantly enhanced activity toward acetyl-lysine.

**Figure 3:**
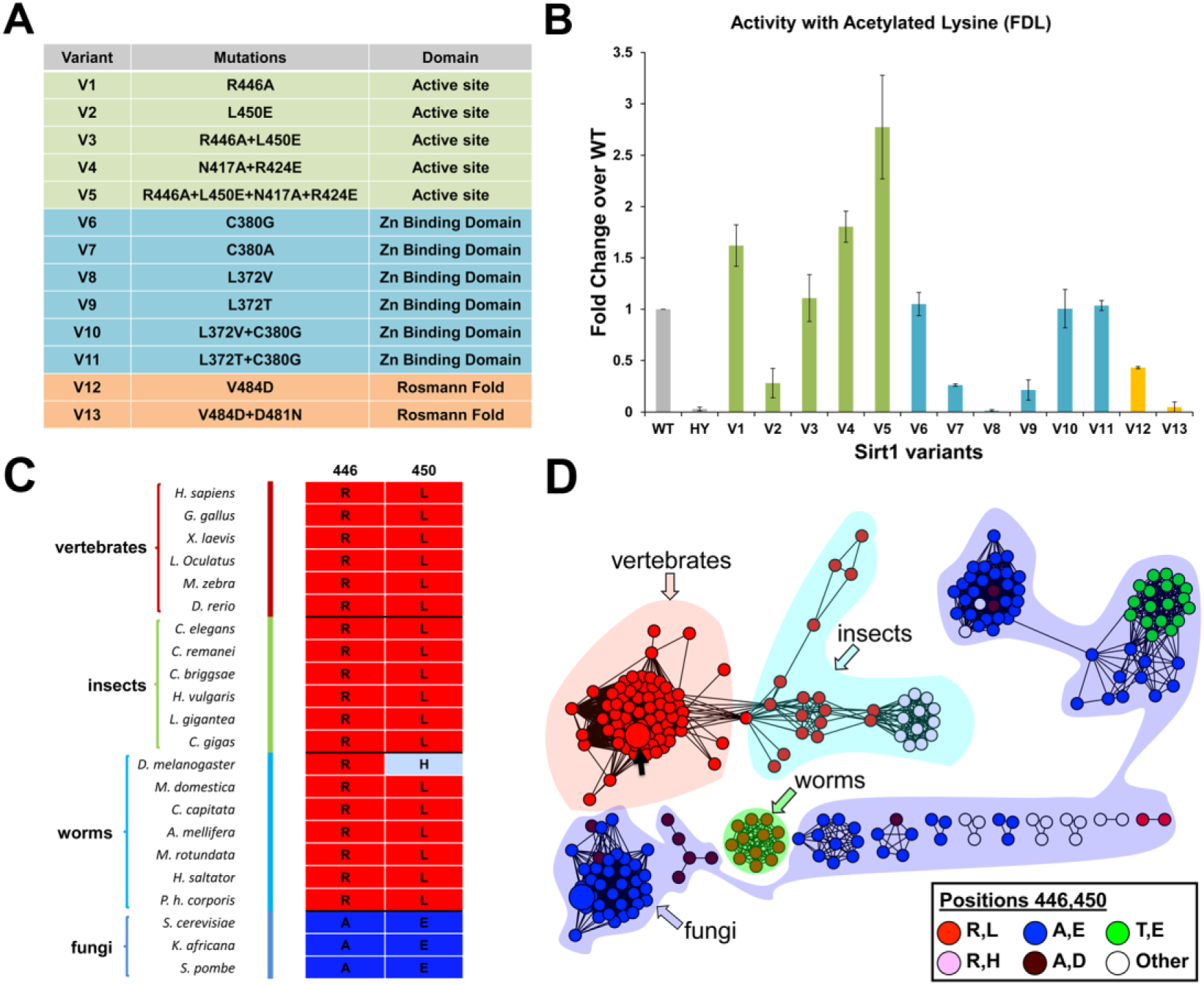
The analysis of 13 hSIRT1 mutants revealing the intramolecular coevolution in hSIRT1. (A) A list of the 13 hSIRT1 mutants generated (V1-V13) classified according to three structural regions in hSIRT1: The active site vicinity (green), Zinc-binding domain (blue) and Rossmann fold (orange) (see also **Fig. 2**). (B) Activity analysis of V1-V13 using the FDL assay performed at 0.9 μM of purified SIRT1 variants. The slopes derived from the FDL experiments were normalized relative to the WT hSIRT1 and results are presented as fold change over the WT (for representative kinetic curves see **Fig. S5**), HY is the inactive hSIRT1 mutant containing the H363Y mutation. Values shown are the mean of three independent repeats while the error bars represent the standard deviation from the mean (C) Amino acid identities of position 446 and 450 in different SIRT1 orthologs highlighting the concerted change from A and E in fungi to R and L in vast majority of all other species shown in the panel (D) An SSN of 103 SIRT1 sequences, as shown in **Fig. 1**, showing the co-evolution between positions 446 and 450 in hSIRT-1 orthologs. Node colors represent hSIRT1 positions 446 and 450 showing the correspondence between the concerted changes in amino acid type of the two positions. The proximity between these positions in hSIRT1 is shown in **Fig. 2**.

Further analysis of other regions in the DAC domain including the Rossmann fold and the Zinc-binding domain (**Fig. 2**) indicates additional intramolecular coevolution in hSIRT1. In the Zinc-binding domain, we assessed the majority of single and double mutants containing the L372T/V and/or C380G/A mutations (V6-V11, **Fig. 3**) to identify a combination of mutations that will not abrogate hSIRT1 deacetylation activity. We found that the L372T and C380G substitutions (V11) that originates from worms SIRT1 orthologs (e.g. *C. elegans*), allows maintaining of deacetylase activity (**Fig. 3**) indicating the coevolution between these residues (**Fig. S6A**). Similarly, only the V484D and D481N double mutant (V13) in the Rossmann fold domain exhibits high deacetylase activity (**Fig. 3**). Structural analysis of positions D481 and V484 shows their spatial proximity suggesting that local excess of a negative charge in the V484D must be compensated by D481N for maintaining hSIRT1 activity (**Fig. S6B**).

### Active site substitutions maintain hSIRT1 activity toward conserved histone substrates

As described above, hSIRT1 V3-V5 mutants contain substitutions for residues found in fungi or fly SIRT1 orthologs and exhibit WT or higher level of activity toward acetylated lysine (**Fig. 3**). To examine V3-V5 activity toward acetylated lysines in universally conserved histones, we first utilized peptides containing H3K9Ac, H3K56Ac and H4K16Ac derived from human histones H3 or H4. The activity of WT hSIRT1 and V3-V5 mutants toward these peptides was measured using ammonia-coupled assay as previously described (Meledin et al., 2013; Smith et al., 2009). This continuous spectroscopic assay, allows the Michaelis Menten (MM) analysis of WT and V3-V5 activity with acetylated peptides to derive the *K*_M_ and *k*_cat_ parameters. Analysis of these variants with the different peptides revealed similar or in some cases even lower *K*_M_ values for V3-V5 relative to the WT (**Fig. 4A-B** and **Figs. S7-8**). Overall, this analysis indicates that the mutations in V3-V5 do not compromise its ability to recognize and catalyze the deacetylation of peptide derived from H3 and H4 histones (for full MM parameters see **Table S2**).

**Figure 4:**
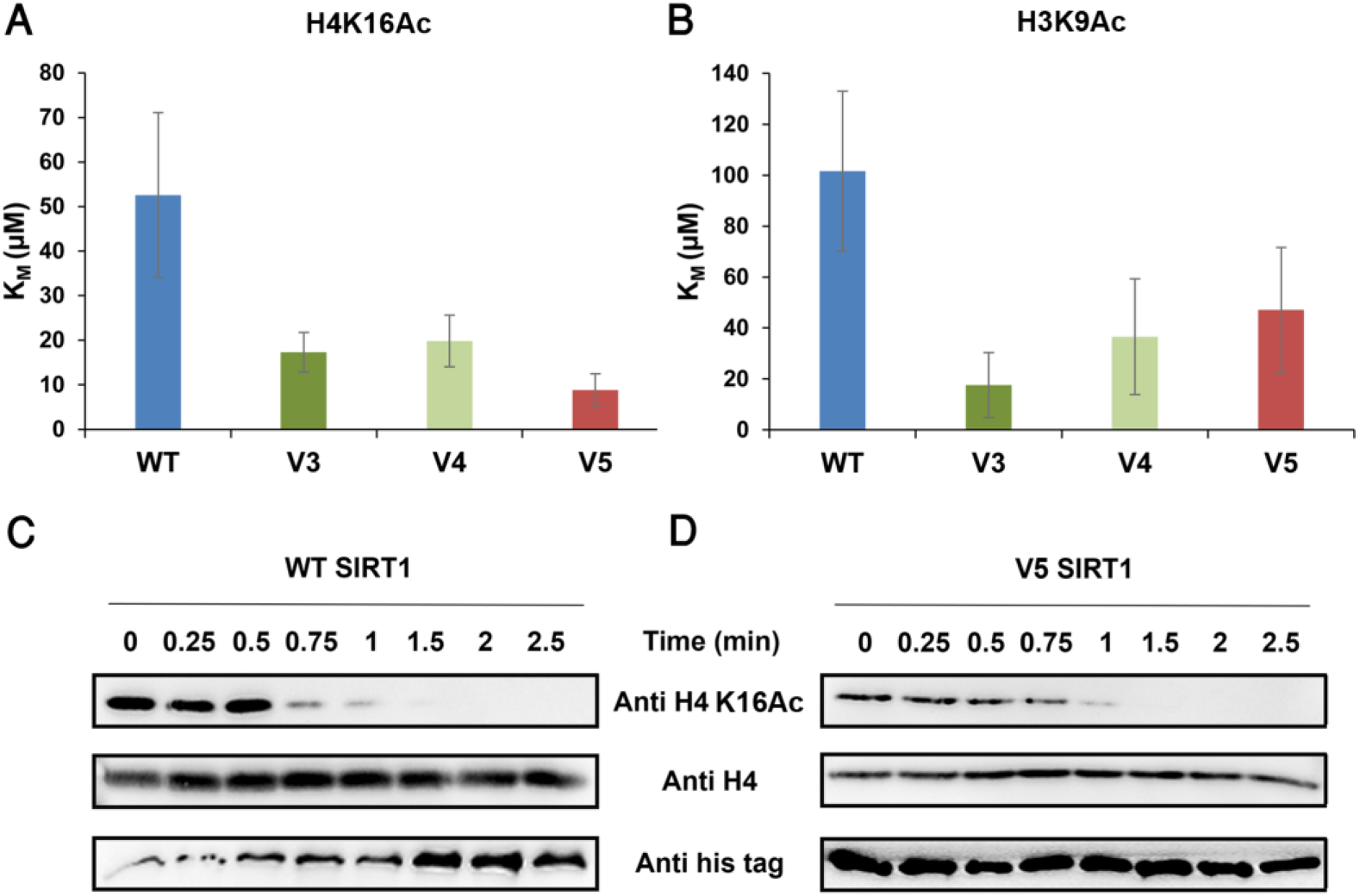
The activity of V3-V5 toward acetylated histones is maintained. (**A-B**) The K_M_ values of the WT and V3-V5 for H4K16Ac and H3K9Ac, respectively, obtained from the MM curves (see **Fig. S7)**. (**C-D**) Western blot kinetic analysis of WT and V5 activity toward H4K16Ac in the context of native histones shows that V5 is fully active toward this substrate. SIRT1 deacetylation activity is detected by the time dependent decrease in H4K16Ac signal. Western blots of WT (**C**) and V5 (**D**) H4K16Ac, H4 and hSIRT1 levels were detected using anti-H4K16Ac antibody, anti-H4 antibody and anti-6xHis antibody, respectively.

To further measure the activity of V3-V5 toward full-length native acetylated histones, we first purified the chromatin fraction from HEK293T mammalian cells. Next, we incubated these variants with the purified chromatin fraction produced from the HEK293T cells for limited time durations. We then utilized western blot analysis to monitor the activity of hSIRT1 variants toward H4K16Ac by following the time dependent decrease in H4K16Ac acetylation signal. In accordance with these results, we found that V3-V5 activity toward H4K16Ac in the chromatin fraction is similar to the WT activity (**Fig. 4** and **Fig. S9**). Overall, these results show that V3-V5 maintain full activity with the conserved acetylated histones.

### Multiple active site substitutions lead to reduced activity toward acetylated p53 and RelA

Our bioinformatics analysis of SIRT1 substrates indicates that while histones are fully conserved, p53 is less conserved and appears only in multicellular organisms (see **Fig. 1** for p53 evolutionary dynamics). We hypothesized that the possibly recent substitutions at positions N417, R424, R446 or L450 (**Fig. 1-2**) may be important for efficient recognition of p53 by hSIRT1. In this case, the V3-V5 mutants, containing two or four active site vicinity mutations originating mainly from Fungal SIRT1 orthologs (**Figs. 1-3**), may exhibit decreased deacetylation activity toward p53. To examine this hypothesis, we first examined V3-V5 deacetylation activity with a K382Ac peptide derived from human p53. We found that the *K*_M_ of V5, containing 4 mutations, toward this peptide is increased by ∼2.5 fold, relative to the WT, indicating a minor decrease in affinity for the V5-p53 peptide (**Fig. 5A, Fig. S8** and **Table S2**).

**Figure 5:**
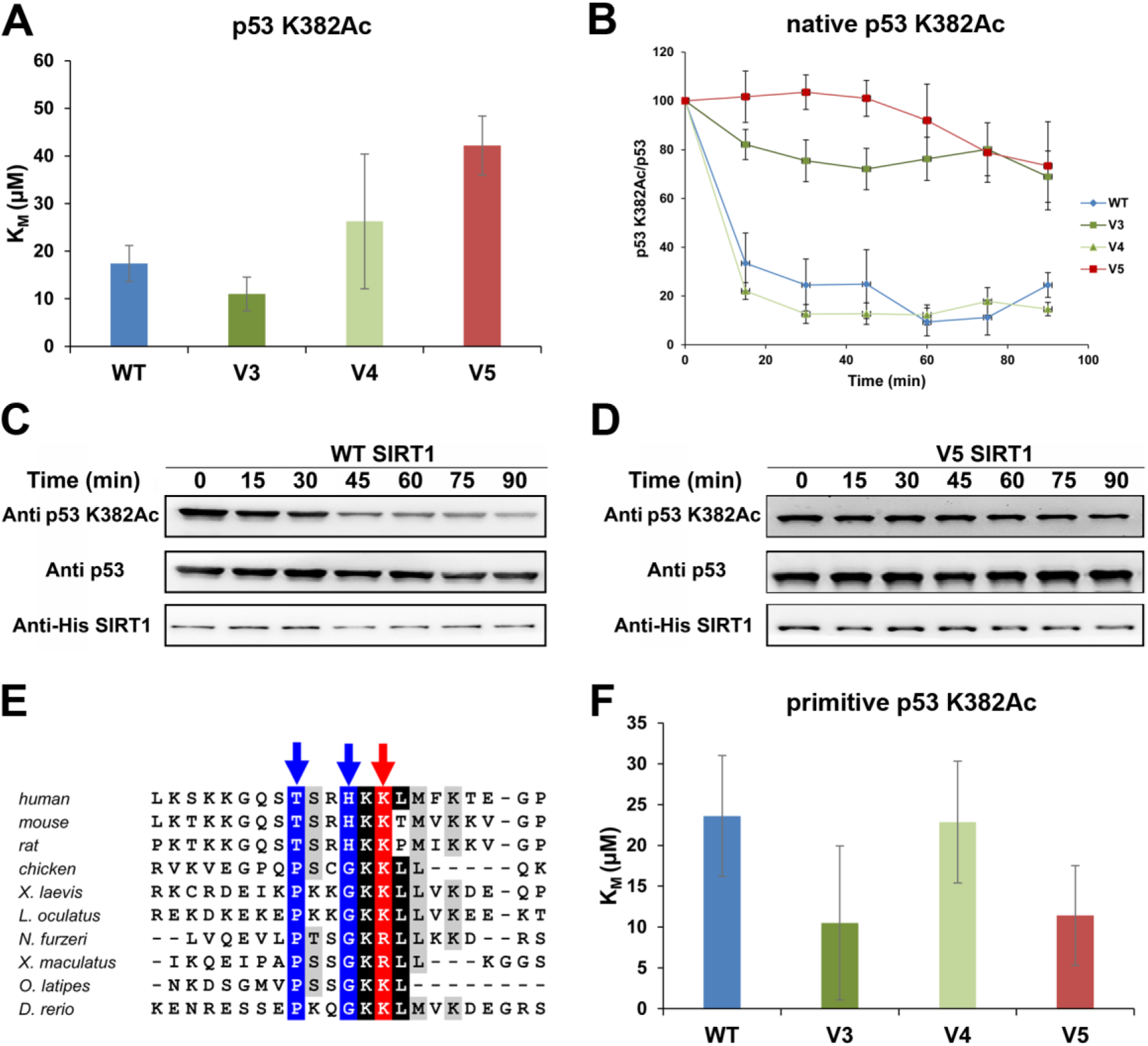
The activity of WT and V3-V5 with acetylated p53. (**A**) The K_M_ values of the WT and V3-V5 with p53-K382Ac peptide obtained from the MM analysis are shown in **Table S2**. (**B**). Western blot kinetic analysis of WT and V3-V5 activity toward p53-K382Ac in the context of native p53 containing AcK at position 382 shows that V3 and V5 exhibit dramatically reduced activity. The time dependent decrease in K382Ac signal was normalized to time 0 and shown as the percentage of p53-K382Ac/p53 bands. Bands intensity were quantified by Image J. Analysis of p53-K382Ac, p53 and hSIRT1 levels were performed using anti-K382Ac antibody, anti-p53 antibody and anti-6xHis antibody, respectively. (**C-D**) Representative blots for WT (**C**) and V5 (**D**). (**E**) A multiple sequence alignment of p53 orthologs around K382 (in human, red arrow). T377 and H380 residues (human p53) are strongly conserved in mammals, but a proline and glycine are present at the same position in other vertebrates. (**F**) The K_M_ values of the WT and V3-V5 mutants with p53-K382Ac primitive peptide obtained from the MM analysis shown in **Fig. S7**.

To further examine V3-V5 activity toward acetylated p53 protein, we used the genetic code expansion approach in *E. coli* to site specifically incorporated *N*-(ε)-acetyl-L-lysine (AcK) (Avrahami et al., 2018; Neumann et al., 2009) into K382 position of p53 during protein expression. The p53 K382Ac was then expressed, purified and utilized as a substrate for the WT and V3-V5 mutants. We used western blot to monitor the time dependent deacetylation activity of hSIRT1 variants with the acetylated p53 protein substrate. Using this assay, we observed a dramatic decrease in V3 and V5 activity toward acetylated p53 protein, relative to the WT, while the activity of V4 was unaffected by the mutations (**Fig. 5B-D** and **Fig. S10**). These results indicate that while V3 and V5 maintain full activity toward histones (**Fig. 4**), their activity toward p53 protein substrate is significantly reduced. In addition, these results indicate that the R446A and L450E mutations are sufficient to dramatically decrease hSIRT1 activity toward p53 protein while the N417A and R424E mutations do not affect hSIRT1 activity toward K382Ac p53. It is also notable that while the activity with the p53 peptide is only marginally affected, the activity with the full protein is considerably more susceptible to the effects of these mutations.

Next, we examined whether natural variations in sequence adjacent to p53 K382 can affect the deacetylation activity of hSIRT1 variants. To explore this possibility, we first performed bioinformatics analysis of p53 orthologs and found that the sequence around K382 contains two conserved substitutions, found in fish and Aves species, at positions −2 and −5 to K382, relative to human p53 (**Fig. 5E**). V3 and V5 contain substitutions that are based on SIRT1 orthologs found in distantly related eukaryotes including yeast (**Fig. 1**) and exhibit lower activity toward human p53. Next, we measured the activity of WT and V3-V5 mutants with p53 acetylated peptide derived from fish or Aves p53 orthologues (**Fig. 5E**). Interestingly, kinetic analysis of V3 and V5 activity with this peptide shows a minor increased *K*_M_ values, relative to the wild-type and V4 proteins (**Fig. 5F** and **Table S2**). These differences in *K*_M_ may indicate that substitutions in the vicinity of p53 acetylation site can contribute to efficient SIRT1-substrate recognition in higher eukaryotes.

Finally, to examine whether V3-V5 variants exhibit altered activity with an additional substrate that appeared later in evolution, we chose to examine SIRT1 activity with RelA/p65 (**Fig. 1**). RelA/p65, belonging to the NFkB complex, was previously shown to be acetylaed at position K310 and is specifically deacetylation by hSIRT1 (Yeung et al., 2004). To examine WT and V3-V5 mutant deacetylation activity with K310Ac peptide derived from RelA/p65, we used the continuous ammonia assay (Smith et al., 2009). We found that all variants exhibit higher *K*_M_ than the WT. V5, containing all 4 active site substitutions, exhibits a significant increase of 10 fold in *K*_M_, relative to the WT (**Table S2** and **Fig. S8**).

### Ancestral sequence reconstruction (ASR) of hSIRT1

Our analysis of V3 and V5 indicates that substitutions of residues near the active site lead to loss of deacetylation activity against p53 protein with no effect on activity with the conserved acetylated histones. To gain more detailed insight into the evolution of this substrate binding region from a common ancestor for the Fungal and Metazoa, we performed ASR analysis of SIRT1 DAC focusing on active site vicinity residues. Given that the co-evolution of amino acid residues is frequently indicative of functional evolution, we first performed residue co-evolution analysis of active site vicinity residues. To perform this analysis, we used more than 6000 sequences of the SIRT1 DAC using the Generative REgularised ModeLs of ProteiNs (GREMLIN) (Balakrishnan et al., 2011) online server. The pairwise coevolution matrix output by GREMLIN was arranged into a network representation using Cytoscape (Shannon et al., 2003). The results were examined to identify any residue networks in SIRT1 that were in close proximity to the substrate binding site. One particular network, consisting of amino acids mutated in V5 (positions 417, 424, 446 and 450) and position 449 was identified with high confidence and was examined in more detail (**Fig. 6**).

**Fig. 6:**
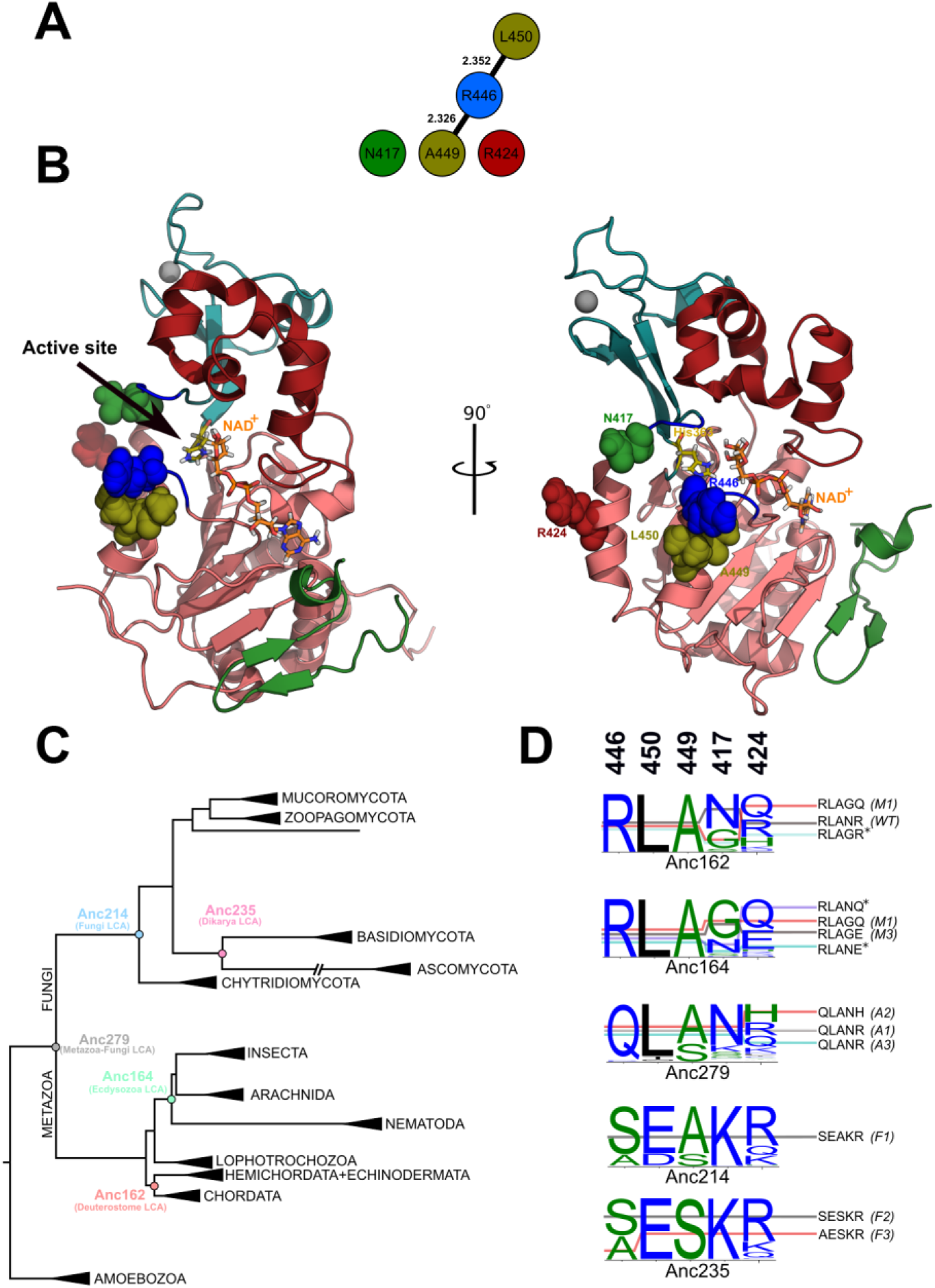
**(A)** Putative specificity-determining positions (SDPs) from the GREMLIN pairwise co-evolution network. Residue positions and identities are labelled according to the hSIRT1 sequence Q96EB6. Edge labels show GREMLIN co-evolution score. **(B)** SDPs shown as spheres on the crystal structure of hSIRT1, 4KXQ. Putative substrate binding/recognition loops are coloured in blue. (**C**) The constrained maximum-likelihood phylogeny of the SIRT1 DAC domain inferred from an alignment of 142 sequences. Key ancestral nodes are indicated. Sub-trees are collapsed and labelled according to taxonomic kingdom (e.g. Amoebozoa), phylum (e.g. Ascomycota), or class (e.g. Insecta). Human SIRT1 is found in the Chordata sub-tree; ySIR2 is found in the Ascomycota sub-tree. (**D**) The posterior probability distribution at the putative specificity determining positions (446, 450, 449, 417 and 424, respectively in the sequence logos) for key ancestral nodes, along with possible combinatorial configurations. Names of hSIRT1 mutants arising from the combinations are shown in brackets. ‘*’ denotes combinations that are not found in the extant sequence set. Sequence logos constructed using WebLogo 3(Crooks et al., 2004).

To examine the evolutionary history of SIRT1, we then used ASR to reconstruct SIRT1 sequences along the evolutionary trajectories of Fungal and Metazoan orthologs. Phylogenetic inference based on an expanded 142-taxon alignment of Metazoan and Fungal SIRT1 sequences was performed using the ML method (see Materials and methods for details, **Fig. 6**). Ancestral sequence reconstruction was performed using the Empirical Bayes (EB) method of Yang in PAML (Yang, 2007), with the LG evolutionary model (Le and Gascuel, 2008) (see Methods). Focusing on the conserved DAC domain of SIRT1 allowed us to obtain high posterior average probabilities for maximum likelihood ancestral sequences.

We expressed and purified an additional series of hSIRT1 variants (M1-3, representing three ancestors on the metazoan branch; F1-3, representing ancestors on the fungal branch; and A1-3, representing common ancestors) containing active site substitutions (at positions 446, 450, 417, 424 and 449) to residue identities that may have been found in ancestral SIRT1 sequences. The ambiguity in the reconstruction at these positions meant that a number of combinations were possible at each ancestral node; we expressed all possible combinations at each node except where analysis of extant sequences indicated that the combination does not exist in extant sequences and is therefore unlikely to have existed in ancestors (owing to coevolution and epistasis) (**Fig. 6A** and **Fig. S11**).

### Experimental analysis of ASR based hSIRT1 mutants

According to the ASR analysis, we generated, expressed and purified seven different hSIRT1 variants with mutations to ancestral states at the key residue positions at the selected ancestral nodes (**Fig. 6**). Previous studies indicated that a shorter version of hSIRT1 containing residues 236-684 (Knyphausen et al., 2016), including the SIRT1 C-terminal regulatory region (CTR), leads to reduced aggregation with complete maintenance of SIRT1 enzymatic activity. Thus, we expressed this N-terminal truncated version of the different ASR based variants and WT for obtaining higher expression level and decreased aggregation relative to the full length hSIRT1. To verify that N-terminal truncation doesn’t affect the specificity toward acetylated p53, we examined the truncated versions of WT and V5 with comparison to the respective full length proteins. We found that the truncations don’t change the activity toward p53 acetylated protein, verifying that the N-terminal region is not associated with p53 recognition (**Fig. S12**).

To examine the activity of the different variants, we first used the FDL assay using the Boc-AcK-AMC substrate, as described above, verifying that all variants were active (**Fig. 6** and **Fig. S13**). Next, we used western blotting as described above, to examine the activity of the different variants against H4K16Ac and p53 K382Ac using native histones extracted from cells and purified acetylated p53, respectively. Interestingly, we found that the A1-A3 variants, corresponding to possible residue configuration of the Metazoa-Fungi common ancestor, exhibited similar activity with these substrates as WT hSIRT1 (**Fig. 7** and **Fig. S14**). In contrast we found that the F1 and F3 variants, corresponding to possible residue configurations of the common ancestor of all Fungi and the common ancestor of Dikarya (yeasts and mushrooms), respectively, maintained activity with the conserved H4K16Ac substrate but exhibit dramatic loss of activity with the p53 K382Ac substrate (**Fig. 7** and **Fig. S14**). This result indicates that while the residue configurations of the Metazoa-Fungi common ancestor were able to catalyse p53 deacetylation, this ability was lost during the Fungal SIRT1 evolution. Finally, the analysis of M1 and M3 variants, generated based on the ancestral sequences from the Metazoa, show that these variants exhibit similar activity as WT hSIRT1 with p53. Overall, these results suggest that the active site configuration of the Metazoa-Fungi common ancestor exhibits multi-specificity towards both histone and p53 substrates, and that this trait was maintained during Metazoan SIRT1 evolution but lost during Fungal SIRT1 evolution.

**Fig. 7:**
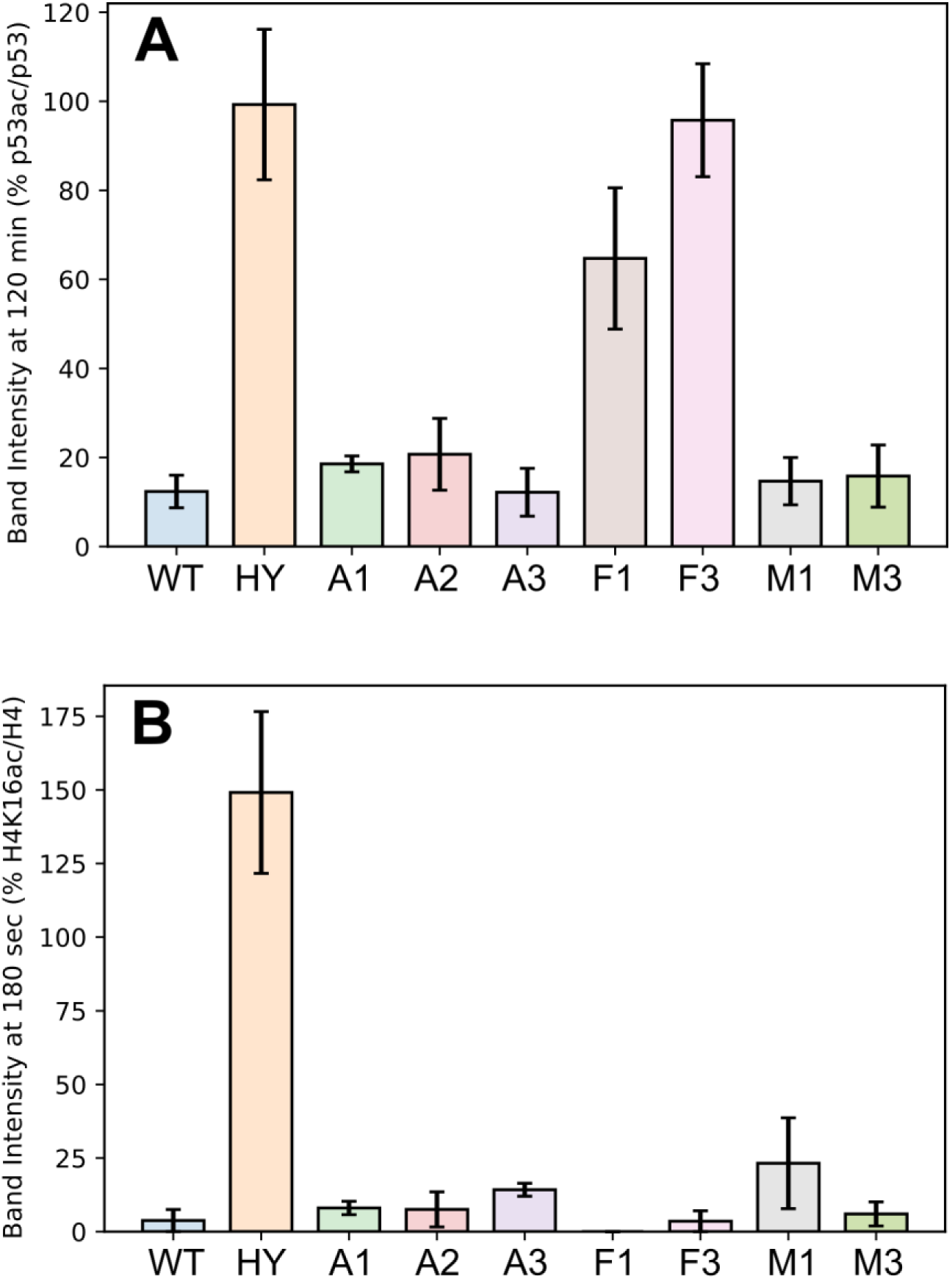
Quantification of western-blot activity assay for the different hSIRT1 ASR based variants with native p53 K382Ac and native H4K16Ac. **(A)** Data for p53 K382Ac was quantified by the band intensity following membrane exposure to anti-p53Ac antibody at 120 minute time point as a percentage of the intensity of the same time point probed with anti-p53, normalized to the percentage of p53 K382Ac/p53 band intensities at t = 0. (**B**) Data for H4K16Ac is quantified by the band intensity following membrane exposure to anti-H4K16Ac antibody at 180 second time point as a percentage of the intensity of the same time point probed with anti-H4, normalized to the percentage of H4K16Ac/H4 band intensities at t = 0. The hSIRT1 HY mutant, containing the catalytic H363Y mutations, served as the negative control. Error bars show the standard error from the mean (SEM) for a minimum of 2 experimental repeats. Representative western blots for all variants are shown in **Fig. S12**.

The lack of activity of V3 with p53, containing R446A and L450E mutations (**Fig. 3**), reveals that the identity of these residues plays a critical role in p53 K382Ac recognition. Similarly, F1 and F3 exhibit complete loss of activity toward p53. Analysis of the substitutions of 446 and 450 in these variants reveals that both variants contains the L450E mutation while R446S and R446A mutations are present in F1 and F3 variant, respectively. These results are in good agreement with the mutations in V3 and reveal L450E as the key specificity determining residue that must coevolve with R446 substitution to maintain high deacetylation activity with acetyl-lysine but reduced activity with p53 (**Figs. 4-7**). Finally, A446 and E450 are conserved in most fungi species (**Fig. 1** and **Fig. S2**) suggesting that Fungal SIRT1 orthologs lost p53 recognition during evolution.

## Discussion

Many enzymes that catalyze the addition or removal of diverse types of PTMs on different protein targets are multi-specific. This phenomenon is extremely hard to decipher and remains enigmatic for many such enzymes (Beltrao et al., 2013). Our study on hSIRT1, as a prominent example, addresses fundamental questions regarding the molecular basis and evolution of multi-specificity in these enzymes. In eukaryotes, enzyme-substrate networks can dramatically expand through evolution upon transition from primitive eukaryotes such as fungi to complex eukaryotes such as humans. Thus, natural changes in enzyme sequences can be related to their growing ability to recognize multiple substrates. Here, we examined this hypothesis by using a bioinformatics-experimental approach to identify and test natural substitutions in SIRT1 taking place during the evolution of eukaryotes. We mainly focused on substitutions of active-site vicinity residues in hSIRT1 and examined the activity of different hSIRT1 mutants with conserved and non-conserved substrates.

hSIRT1 is an excellent model system due to its deacetylation activity toward a large number of acetylated residues in proteins and its involvement in the regulation of a variety of cellular processes in health and disease (Lavu et al., 2008). A recent study that examined the deacetylation profile of all human SIRT1-7 enzyme with 6802 different acetylated peptides, representing the human acetylome, revealed that thousands of acetyl-lysine sites are targeted by these enzymes (Rauh et al., 2013). In addition, other studies revealed that the substrate preference of sirtuins is not limited to acetyl-lysine residues but these enzymes, primarily SIRT6, can recognize different fatty-acid acyl-lysine modified substrates (Feldman et al., 2013; Jiang et al., 2013). However, despite extensive studies, little is known regarding the molecular basis for the high degree of sirtuins multi-specificity. Several studies on the specificity of hSIRT1, using different types of peptide arrays (Blander et al., 2005; Garske and Denu, 2006), revealed no clear consensus sequence for acetyl-lysine recognition. In contrast to the multi-specificity of hSIRT1, the substrate repertoire analysis of the ySir2 ortholog from *S. cerevisiae* revealed its narrow specificity toward conserved histone acetyl-lysine residues (Bheda et al., 2012). This observation suggests that multi-specificity in SIRT1 can be acquired or lost through evolution, nevertheless, its molecular basis is still unknown.

Our bioinformatics approaches for the analysis of SIRT1 sequences, first allowed the identification of partially conserved residues in SIRT1 that coevolve (**Fig. 1-3**). Previously, intramolecular co-evolution was shown to be an extremely powerful tool for revealing contact constrains within proteins allowing the elucidation of the three dimensional structure of many proteins (Marks et al., 2012; Ovchinnikov et al., 2017, 2015). Indeed, experimental analysis of several hSIRT1 mutations to residues, found in SIRT1 orthologs, showed that combination of mutations is needed for maintaining hSIRT1 deacetylation activity (**Fig. 3**). These results suggest that intramolecular coevolution in SIRT1 plays an important role in maintaining its catalytic activity. Our findings further support the notion that co-evolving residues are highly important for maintaining protein function through evolution.

The identification of partially conserved residues at the vicinity of the hSIRT1 active site allowed us to generate and focus on the V3-V5 variants, containing 2 and 4 substitutions at the vicinity of hSIRT1’s active site that originate from SIRT1 orthologs (**Fig. 1** and **Fig. S2**). Our findings that V3 and V5 exhibit high activity toward conserved histone substrates but a dramatic loss of activity toward acetylated p53 (**Figs. 4-5**) suggest that the identity of residues in positions 446 and 450 plays a key role in dictating p53 deacetylation by SIRT1 (**Fig. 1**). ASR followed by experimental examination of SIRT1 variants containing ancestral substitutions at active-site vicinity residues reveal the dynamics of p53 recognition through evolution. These results indicate that the Fungal-Metazoa SIRT1 common ancestor had some activity with multiple substrates, and that evolution of true multi-specificity requires continual selective pressure via the presence of certain substrates. For example, the absence of p53 in Fungi alleviates selective pressure to maintain activity against p53 and this function is lost due to neutral drift or mutations; in Metazoa there has been selective pressure to maintain activity on p53 and this is primarily driven by the coevolving regions we identified in this study.

Previously, the function and evolution of promiscuous enzymes that catalyse the chemical transformation of native and non-native substrates were extensively studied (Babtie et al., 2010; Tawfik and Tawfik, 2010). Directed evolution of these enzymes revealed the robustness of the native enzyme activity to mutations that dramatically affect the promiscuous non-native enzyme activities (Aharoni et al., 2005). In analogy with these studies, it is tempting to speculate that the activity of multispecific enzymes such as SIRT1 toward conserved substrates (e.g. histones) is robust to natural substitutions leading to reduced activity with modern substrates (e.g. p53). However, additional studies with other multi-specific enzymes are required to thoroughly examine this hypothesis.

Overall, our study reveals residues that are critical for SIRT1 multi-specificity and sets the stage for additional analysis of SIRT1 substrates to map in details the evolutionary dynamics of SIRT1 multi-specificity. In addition, future work should reveal the structural basis for multi-specificity and SIRT1 by careful analysis of the effect of mutations identified in this study on SIRT1 structure and substrate recognition. Finally, our approach can be further utilized for the examination of a variety of other multi-specific PTM catalyzing enzymes including kinases/phosphatases, methyl-transferases/demethylases and ubiquitin-ligases/deubiquitinases to study their molecular basis for multi-specificity and shed new light on the evolution of enzyme-substrate recognition in these diverse and important systems.

## Material and Methods

### Collecting sirtuin sequences and generating a sirtuin-superfamily wide sequence similarity network

The UniProtKB(UniProt Consortium, 2013) and NCBI(NCBI Resource Coordinators et al., 2018) sequence databases were searched for hits of sequence patterns associated with sirtuin proteins. Signatures from InterPro (Finn et al., 2017) (IPR026590, IPR026591, IPR003000, IPR017328, IPR026587, IPR027546 and IPR028628) and Pfam (Punta et al., 2012) (PF02146, PF13289) yielded 10,273 unique sequences with pattern matches. The Structure-Function Linkage database tools (Akiva et al., 2014; Barber and Babbitt, 2012) were then used to generate a representative SSN, as described before (Akiva et al., 2017; Atkinson et al., 2009). In this network, nodes represent sets of proteins that share >60% sequence identity as measured by Cd-hit (Li and Godzik, 2006), and edges represent a mean *E*-value more significant than 1×10^−18^ between all pairwise *E*-value scores calculated between the sequences represented in each node. Identifying this cutoff was obtained by manual sampling of several edge inclusion thresholds until a reasonable reconciliation was achieved between distinct similarity clusters and representation of remote similarities between them. Networks in this paper are visualized by Cytoscape (Shannon et al., 2003) (organic layout).

### Phylogenetic profiles of hSIRT1 Substrates

We complied a set of organism that sample the phylogenetic tree of eukaryotes and have fully-sequenced genomes. For each organism, we combined EggNog (Huerta-Cepas et al., 2016) and Inparanoid (Sonnhammer and Östlund, 2015) to find orthologs of hSIRT1 substrates. These auto-generated lists of orthologs were further validated by finding best BLAST (Altschul et al., 1990) reciprocal hits between the human proteome and any of the eukaryotic proteomes (downloaded from UniProtKB (UniProt Consortium, 2013) and EnsEMBL (Zerbino et al., 2018)). These results were manually examined and then cross-validated with literature-documented phylogenetic models for each substrate, whenever available. Fig. 1 includes a color-coded summary of the results: From white (no ortholog was detected) to dark red (highly conserved ortholog). Color intensity is proportional to the bit-score (as computed by BLAST), divided by the alignment length. To evaluate the conservation of putative acetyl-Lysine residues in the substrates, we aligned orthologs of each substrate using MAFFT (Katoh, 2002) and focused on the segments that align with the human acetyl-Lysine. Fig. 1 summarizes the results of these analysis by the letter “K” for each substrate and organism. Absence of the letter means that no Lysine was found in the relevant aligned segment.

### Generation of SIRT1 mutant plasmids

The p38 plasmid containing hSIRT1 (UniProt Q96EB6) gene fused N-terminal 6xHis tag was obtained as a kind gift from Haim Cohen lab, the Bar-Ilan University, Israel. The p38 was modified by deleting one of the two KpnI restriction sites to generate the p38d plasmid. All further genetic manipulations were based on the p38d plasmid. For generating hSIRT1 mutants in the DAC domain, the mutants were cloned into p38d using KpnI and HindIII. The DAC domain mutants were obtained by a PCR reaction in several steps. The First step was performed to amplify the DAC domain in two fragments with a primer that includes the relevant mutation. The two fragments were purified using the GeneJET PCR extraction kit (Thermo), followed by assembly PCR and further amplification of the whole mutant DAC domain. The KOD polymerase (Merck) was used for all PCR amplifications. The resulting mutated DAC PCR fragments were incubated with DpnI (Thermo) for 30 min at 37°C followed by 10 min of heat inactivation at 80°C. The DAC mutants were further purified and cloned to the digested p38d vector using NEB builder Gibson mix (NEB). The V5 mutant was ordered as a synthetic gene, amplified by PCR and cloned as described above. The final product was transformed to NEB competent bacterial cells and plated on LB agar containing 100 µg/ml Ampicillin. Positive colonies were verified by PCR with dreamtaq polymerase (Thermo), followed by purification using NucleoSpin Plasmid EasyPure kit (Macherey-Nagel) and sequenced to verify the correct mutations.

### Expression and purification of SIRT1 mutants and PNC1 in *E. coli*

Plasmids containing the WT and mutant hSIRT1 genes were transformed into Rosseta 2 *E. coli* competent cells, grown in TB medium containing ampicillin and chloramphenicol at 37°C to an OD_600_ of 0.6 and induced with 0.8 mM IPTG followed by 16 hours of incubation at 16°C. Cells were lysed by sonication in lysis buffer containing 40 mM Tris-HCl pH 8, 200 mM NaCl, 10 mM MgCl_2_, 1:2000 EDTA free-protease inhibitor cocktail (Calbiochem) and Benzonaze (Mercury). Cell debris were removed by centrifugation at 10,000g at 4°C for 30 min. The supernatant was loaded onto pre-equilibrated nickel beads, washed with wash buffer containing 40 mM Tris-HCl pH 8, 200 mM NaCl, 10 mM MgCl_2_ and 20 mM imidazole and eluted with elution buffer (similar to wash buffer but containing 500 mM imidazole). The eluted protein was dialyzed with activity buffer containing 25 mM Tris-HCl pH 8, 200 mM NaCl, 1mM MgCl_2_ and 2.5mM DTT, for imidazole removal. The purity of the proteins was assessed by SDS-PAGE on 10% gel and protein concentration was measured by the Bradford method using bovine serum albumin as the standard. The PNC1 plasmid was obtained as a kind gift from Jessica L will and Jorge C. Escalante-Semerena from the University of Georgia and the protein was purified as previously described(Garrity et al., 2007).

### Fluor de lys (FDL) activity assay for WT and SIRT1 mutants

The activities of the mutants were measured by FDL assay using protected acetylated lysine substrate conjugated to a 4-amino-7-methylcoumarin group (AMC) at the carboxyl terminus. The assay was performed using the following protocol: 30 µl of the hSIRT1 WT or mutants at 1.5 µM were added to 20 µl of the reaction mix containing 12.5 mM NAD+, 1.25 mM AcLys-AMC in 50 mM Tris-HCl pH 8.0, 137 mM NaCl, 2.7 mM KCl, 1 mM MgCl_2_ and Bovine Serum Albumin (BSA) 1 mg/ml (FDL buffer). The mixture was incubated in black 96 well plates (Greiner) covered with aluminum foil on ice. The reaction was stopped at different time points by adding 50 µl developer solutions (3 mg/ml Trypsin, 0.05 mM HCl in FDL buffer with BSA). After adding the developer, the plate was incubated in 37°C for 40 min and fluorescence intensities were measured using Infinite M200 plate reader (Tecan).

### Ammonia coupled assay with acetylated peptides

The ammonia coupled assay was performed as previously described(Smith et al., 2009) with minor modifications. The ammonia assay kit (Sigma) and relevant acetylated peptides (Peptron) were used for preparing the reaction mix. The sequences of all peptides used in this study are shown in **Table S3**. This mix contained 75 µl activity buffer, 1 µl of L-Glutamate Dehydrogenase (Ammonia kit, Sigma), 2 µl of 150 mM NAD+, 2 µl of 1 µM purified PNC1, and 16 µl of 1.5 µM hSIRT1. Acetylated peptides were added at different concentration and the volume was completed to 100 µl with DDW. The reactions were measured at 320 nm for 30 min at 30°C using Infinite M200 plate reader.

### Chromatin fractionation and histone deacetylation assay

Chromatin fractionation and deacetylation assay on histones was performed as previously described (Gertman et al., 2018). The deacetylation assay was performed using 70 µl of chromatin fraction, 20 µl 125 mM NAD+, 30 µl of activity buffer (30 mM Tris-HCl pH 8, 4 Mm MgCl_2_ and 1 mM DTT) in the presence of 40 µl of 1.5µM purified hSIRT1. The reaction was stopped at different time point of 0, 15 sec, 30 sec, 45 sec, 1 min, 1.5 min, 2 min and 2.5 min. To detect the decrease in acetylation of H4K16Ac, the samples were resolved on 15% SDS-PAGE gel and transferred with Tran-Blot Turbo Transfer Pack (Bio Rad) using Trans Blot Turbo Transfer System. The membrane was then probed with several antibodies: Rabbit α-H4 (ab10158), Rabbit α-H4K16 (ab109463) (abcam) followed by a secondary antibody goat α-rabbit. For the analysis of hSIRT1 levels, α-his tag conjugated to HRP was used. Western blot bands were quantified using Image J program.

### Purification and deacetylation assay on native p53

The full length proteins were expressed in bacteria that incorporated the non-natural amino acid (acK) by expending the codon usage. The p53 plasmid with the stop codon in the position 382 was cloned to the pCDF Duet plasmid and purified as previously described (Arbely et al., 2011). In the deacetylation assay we used 140µl of 4µM of the full length substrate, 80µl of 0.8µM of purified hSIRT1 or mutants and 100µl deacetylation buffer (25 mM Tris, pH 8.0, 137 mM NaCl, 2.7 mM KCl, 1 mM MgCl_2_, 1 mM NAD+) (Knyphausen et al., 2016). The reaction was the incubated at room temperature and samples were removed at different time points of 0 min, 15 min, 30 min, 45 min, 60 min, 75 min, 90 min and 120 min. All the samples were analysed by western blot with several antibodies: Rabbit α-p53 (#9282), Rabbit α-p53 ac-K382 (#2525) (Cell Signaling) followed by goat α-rabbit or α-mouse antibody conjugated to HRP. Analysis of hSIRT1 levels and band quantification were performed as described above.

### Coevolution analysis, maximum-likelihood phylogenetic inference and ancestral sequence reconstruction

Coevolution analysis was performed on an automatically compiled >6000 sequence MSA of the SIRT1 DAC (using the hSIRT-1 DAC sequence as the initial search sequence) using the Generative REgularised ModeLs of ProteiNs (GREMLIN) (Balakrishnan et al., 2011) online server (http://gremlin.bakerlab.org/). The pairwise coevolution matrix output by GREMLIN was arranged into a more informative network representation using Cytoscape(Shannon et al., 2003), with an edge cut-off at a GREMLIN raw co-evolution score of ∼1.28 to obtain well-delineated and mutationally viable modules of co-evolving residues.

For construction of a maximum likelihood tree, an EFI-EST enzyme similarity network (Gerlt et al., 2015) were used to collect sequences orthologous to hSIRT1, using the *Xenopus laevis* SIRT1 sequence as seed. Using incremental percent identity edge cutoffs in Cytoscape (Shannon et al., 2003), clusters delineated by sirtuin subfamily were obtained. The SIRT1 cluster was identified by locating the human SIRT1 sequence (Q96EB6). We explored this cluster to ensure that the sequences found were orthologous to SIRT1.

Thus, ∼ 400 sequences of hSIRT1 likely orthologues were collected from a number of Eukaryotic kingdoms. Incomplete fragment sequences were purged by manual inspection. Sequences with greater than 90% sequence homology were clustered and represented by a single sequence with CD-HIT (Li and Godzik, 2006) thereby removing redundant sequences from the sequence set. The PROMALS3D alignment program (Pei et al., 2008) was used to align these sequences. The alignment was extensively inspected and the highly divergent N- and C- terminal domains removed. Another round of CD-HIT was followed by PROMALS3D alignment on the truncated sequences. Gaps and alignment errors were corrected through careful manual inspection of the resultant alignment. In particular, highly divergent and un-alignable insertions were observed in most non-Chordate phyla in the zinc-binding domain; these were removed except in the Chordates, where they were deemed to be truly homologous under parsimony criteria. The sequence set was manually enriched with chordate sequences using profile alignment with HMMer (Johnson et al., 2010). Alignments were benchmarked *via* structural alignment of the orthologs ScSIR2 (PDB:4IAO) and hSIRT1 (PDB:4KXQ;4IG9).

Phylogenies were calculated using IQ-TREE (Nguyen et al., 2015) a final alignment of 142 sequences, using ModelFinder (Kalyaanamoorthy et al., 2017) to find the best-fit evolutionary model and the UltraFast (Minh et al., 2013) bootstrap method to evaluate non-parametric bootstrap branch supports using 1000 pseudoreplicate trees. For inference of constrained-topology phylogenies, the constraint tree topology was manually constructed in accord with the prevailing species trees for the relevant taxa (dos Reis et al., 2015; Spatafora et al., 2016). All statistical topology tests were performed in IQ-TREE using 10,000 pseudoreplicate trees. Ancestral sequence reconstruction was performed using the CodeML program in PAML (Yang, 2007), using the LG evolutionary model (Le and Gascuel, 2008).

## Acknowledgements

We thank Danny Tawfik for helpful discussions and Michael Lammers for sending plasmids.

## Supplementary Information

**Fig. S1:**
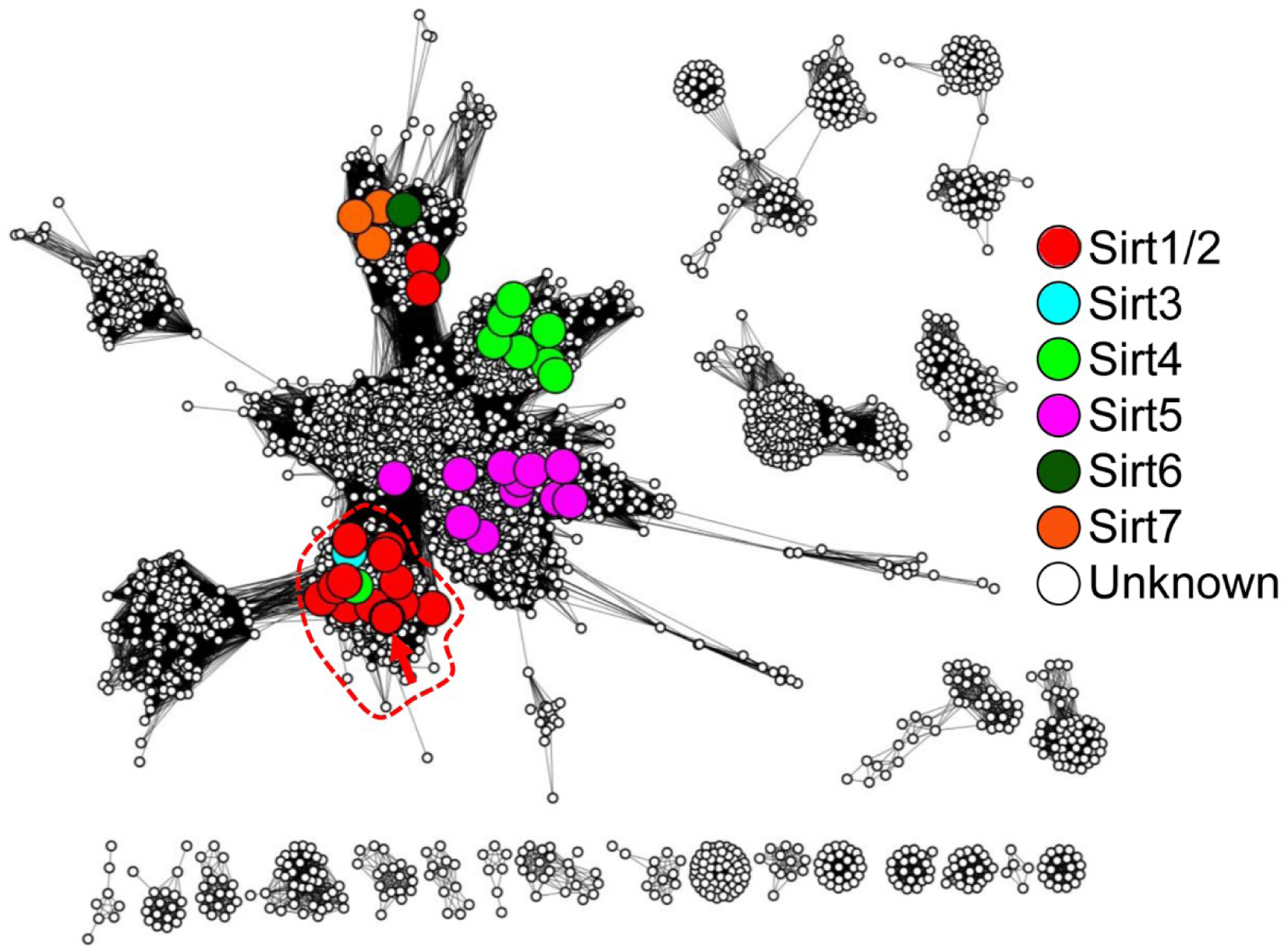
A representative sequence similarity network of the Sirtuin superfamily. Nodes represent proteins that share 80% sequence identity, and edges represent average *E*-Value score below 1*10^−35^. Colored, large nodes represent sequence sets that include at least one experimentally-verified member of the Sirtuin superfamily. In the specific *E*-value used here, very clear separation between large cluster (that represent sequences that are more similar to each other to any other sequences) is observed. The non-colored nodes represent uncharacterized sirtuins, hinting at yet unexplored sequence space of the superfamily. A red arrow points at hSIRT-1. Note that Sirt1/2 are colored the same, to clearly resolve the confusing, yet widely used naming conventions for these proteins. The upper cluster include “Sirtuin-1” superfamily members that belong to plants; this exemplifies, again, the confusing naming standards for sirtuins.

**Fig. S2:**
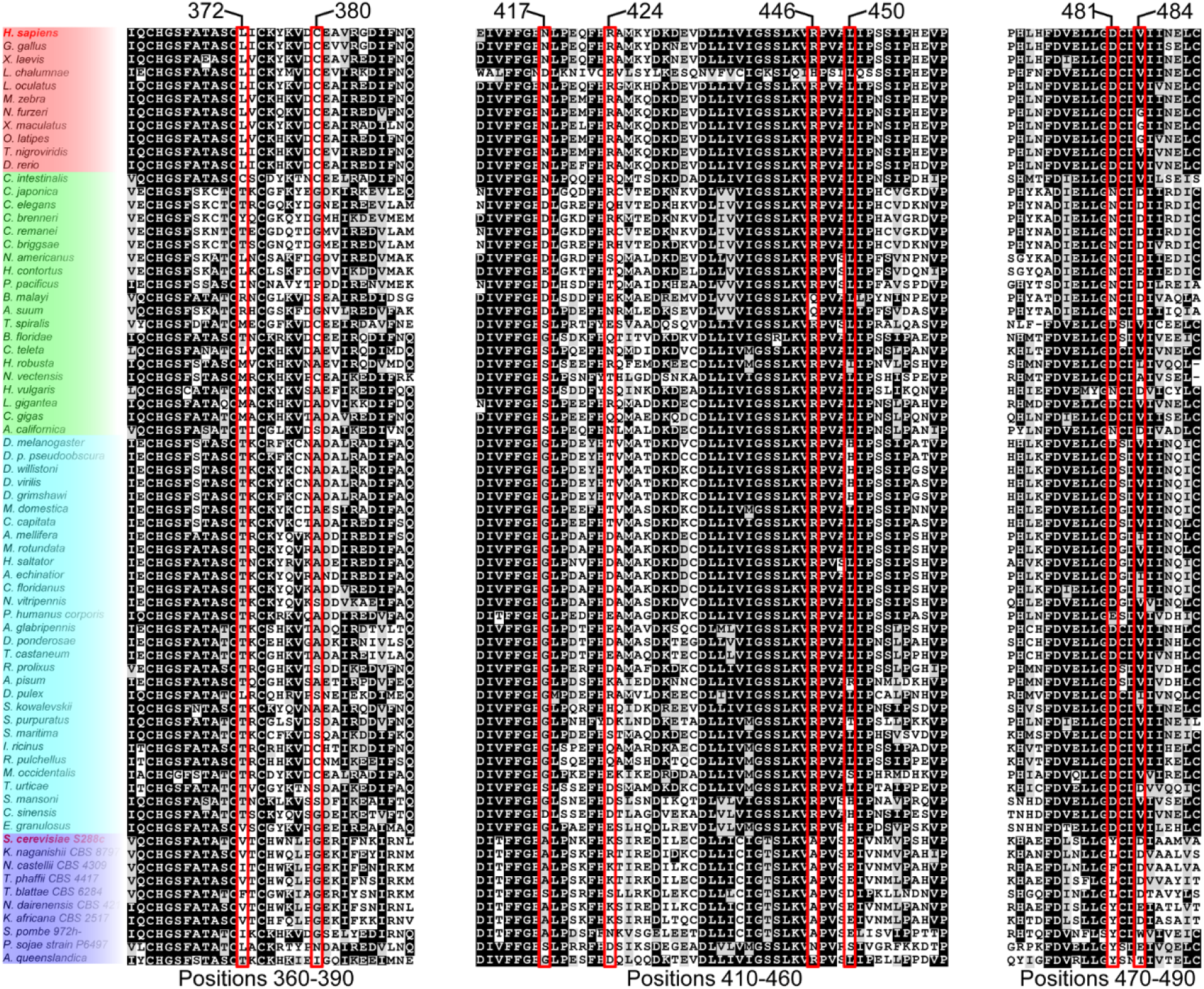
Multiple sequence alignment of SIRT1 likely orthologs. The segments containing the studied substitutions are shown, where the positions are in red boxes. The overall high level conservation of the DAC domain is evident. Organisms’ background colors are grouped into vertebrates (red), invertebrates I (green, mainly mullusca and nematodes), invertebrates II (cyan, mainly Ecdysozoa [insects]) and fungi (blue).

**Fig. S3:**
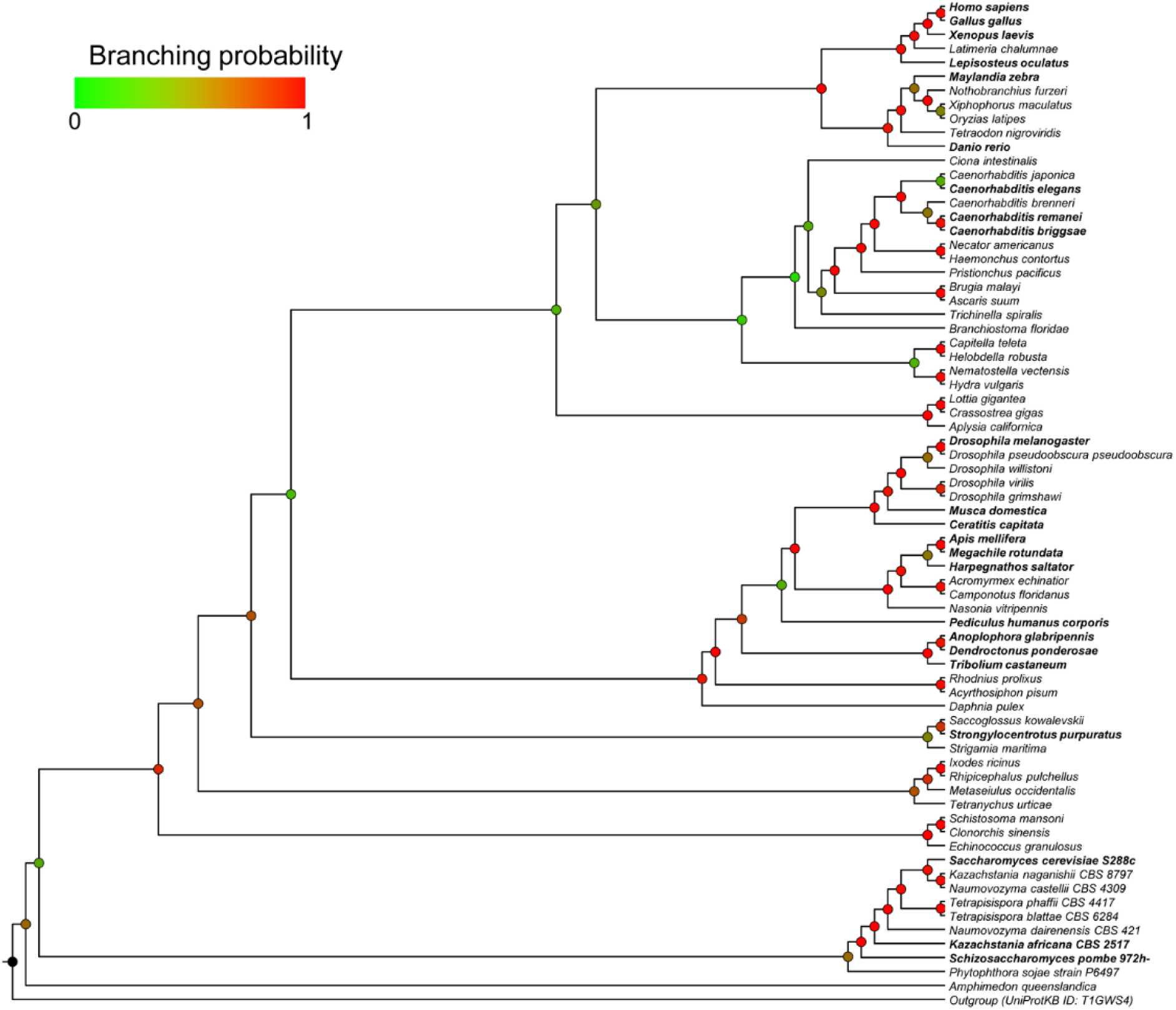
Representative tree of eukaryotic orthologs of SIRT1. The phylogenetic tree, generated using MrBayes, includes 72 of the 103 sequences of SIRT1 orthologs. In bold are organism names that appear in **Fig. 1**.

**Fig. S4:**
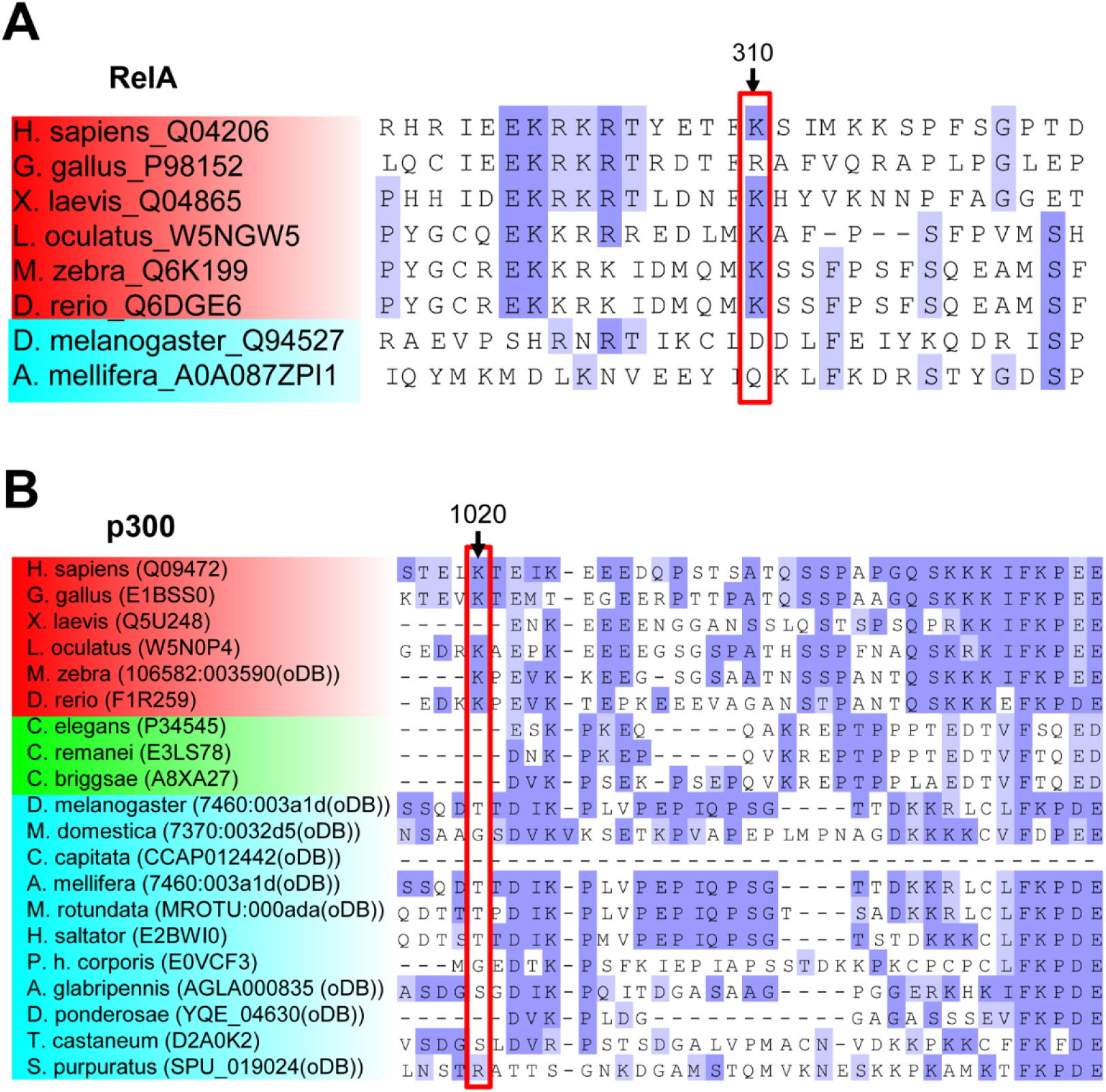
Sequence alignment of RelA/p65 and p300 orthologs emphasizing the conservation of position 310 and 1020 in human RelA and human p300, respectively. Orthologs are based on consensus between diverse databases (OrthoDB, InParanoid, EggNog and EnsEMBL) and^1,2^. Protein identifiers are UniProtKB accession numbers or OrthoDB numbers.

**Fig. S5:**
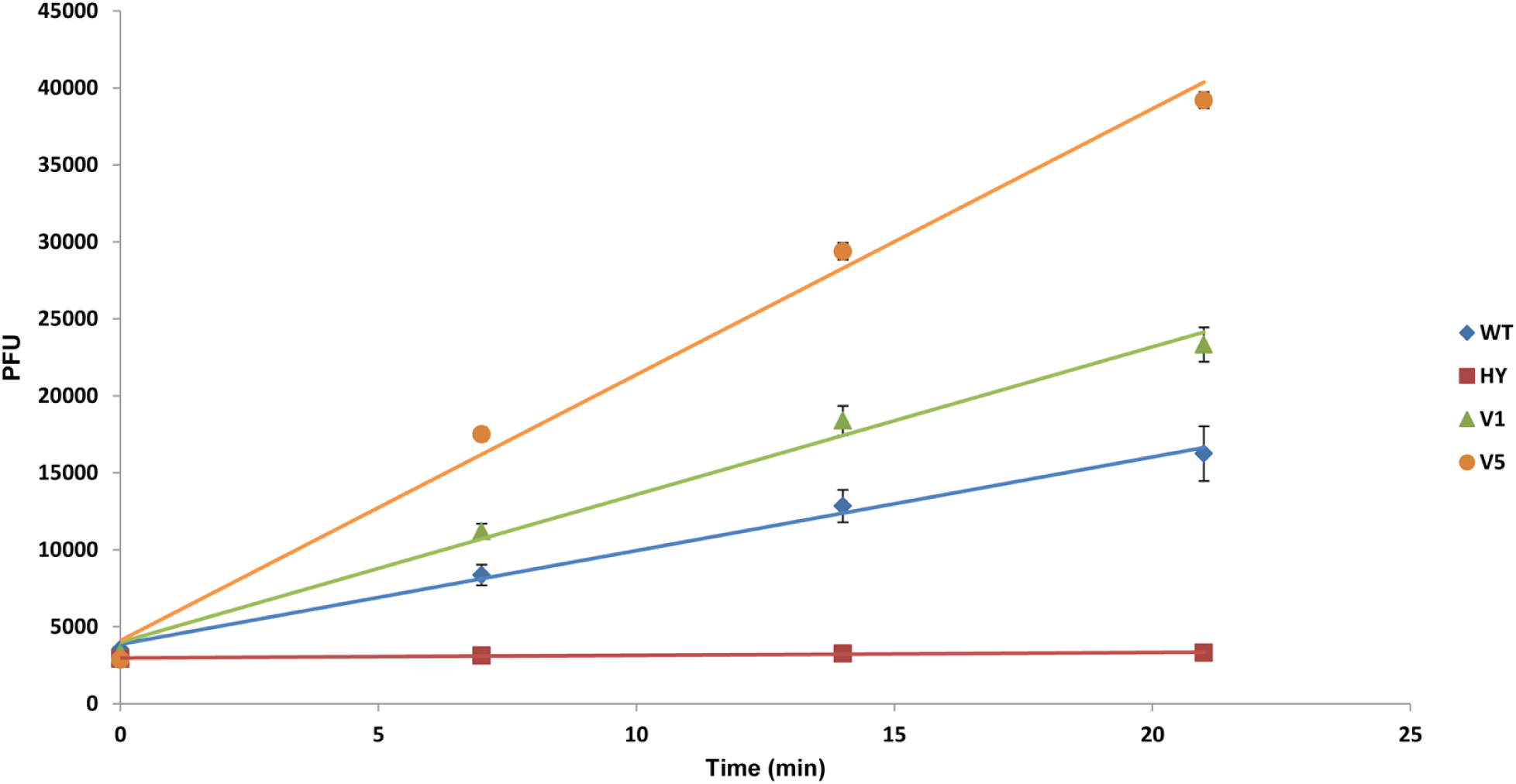
Representative Fluor De lyse (FDL) analysis of hSIRT1 mutants with protected acetyl-lysine-AMC substrate. Fluorescent signal was measured at different time points after stopping the catalytic deacetylation reaction. Initial rates were calculated by fitting the data to linear equation and comparing to the slope obtained for the WT (**Fig. 3B**). The inactive HY hSIRT1 mutant containing the H363Y mutation serves as a control. All measurements were performed in triplicates and the standard deviation from the mean is shown.

**Fig. S6:**
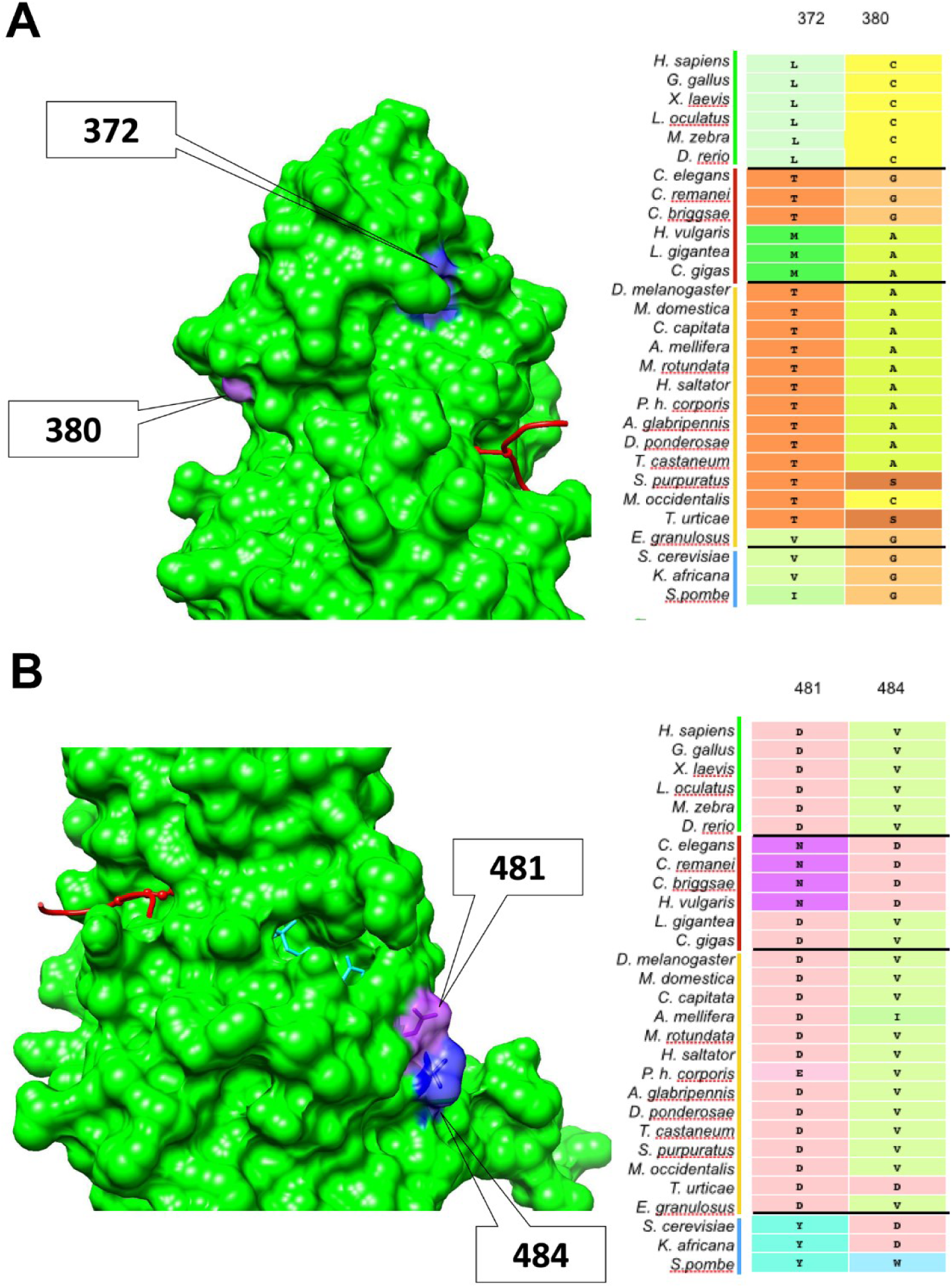
Coevolution analysis of specific positions in SIRT1. (**A**) Analysis of positions 372 and 380 located at the Zinc-binding domain. Left: Surface representation of hSIRT1 structure with positions 372 and 380 highlighted. Right: Sequence alignment of SIRT1 orthologs showing amino-acid identities at these positions. (**B**) Analysis of positions 481 and 484 located at the Rossmann fold domain. Left: Surface representation of hSIRT1 structure with positions 481 and 484 highlighted. Right: Sequence alignment SIRT1 orthologues showing amino-acid identities at these positions.

**Fig. S7:**
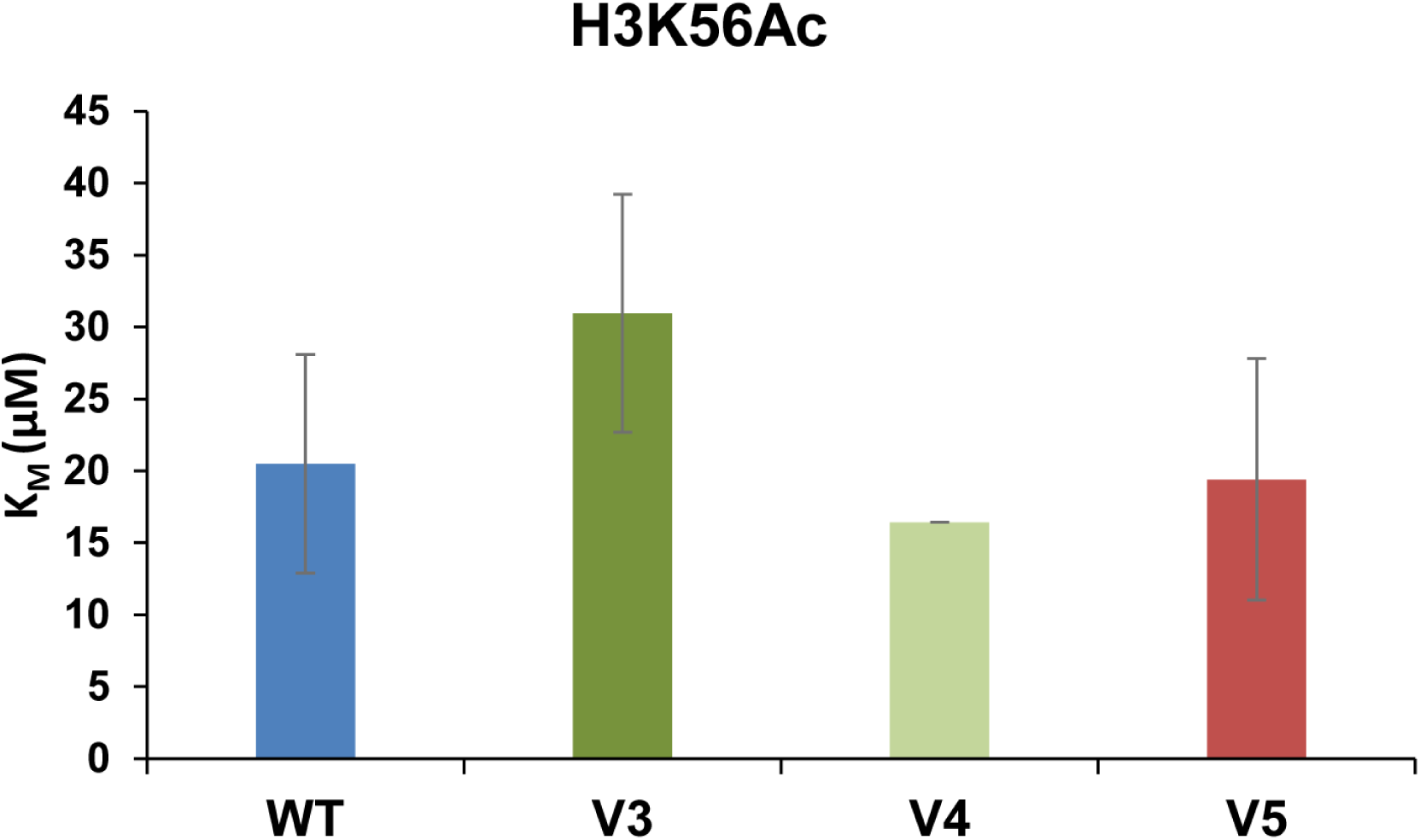
The activity of V3-V5 toward H5K56Ac is maintained. The K_M_ values of the WT and V3-V5 for H3K56Ac measured by the continuous assay^3^ following incubation of the hSIRT1 variants with different concentrations of H3K56Ac peptide. Full MM curves are shown in **Fig. S8**.

**Fig. S8:**
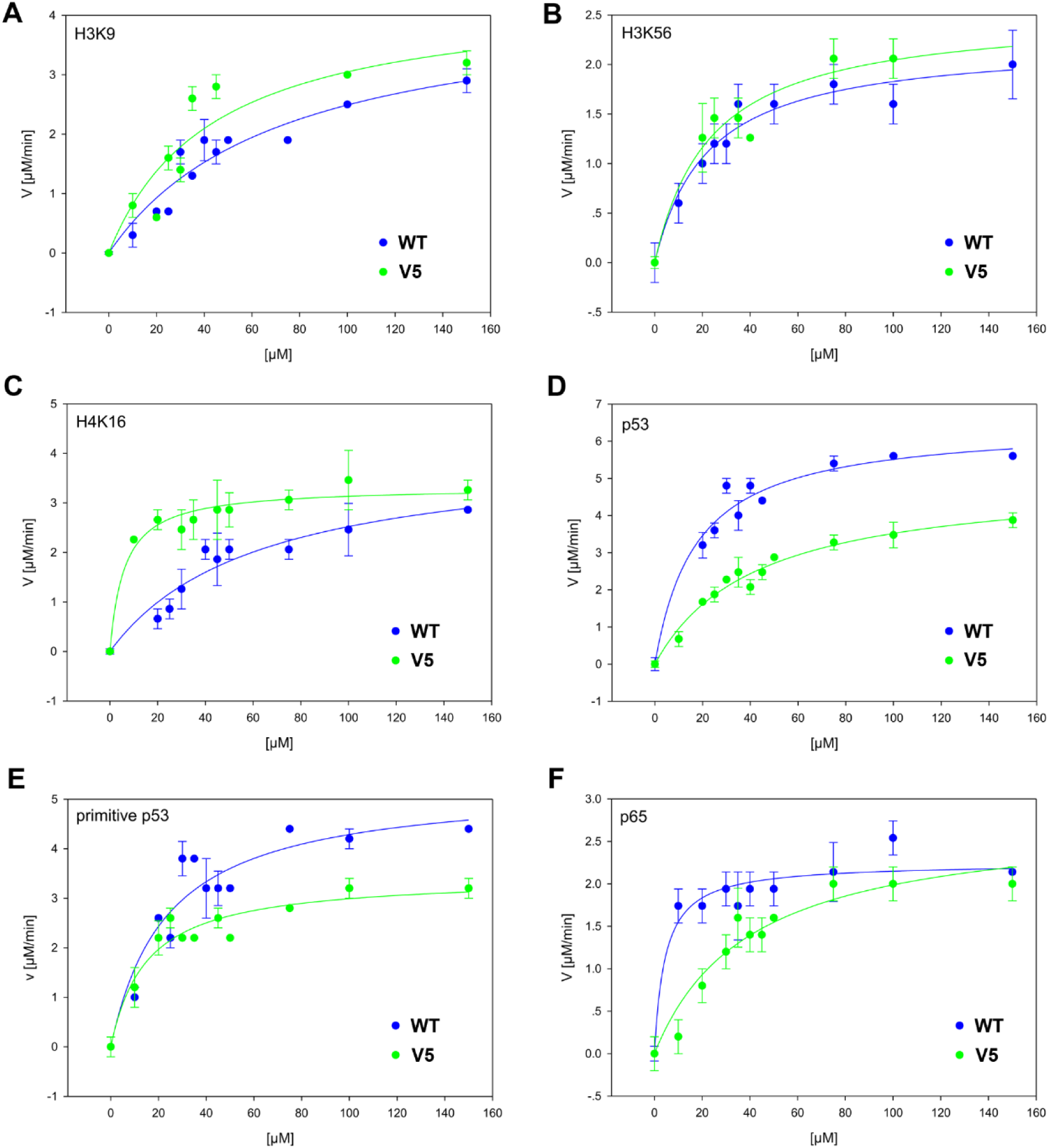
Michaelis Menten (MM) analysis of the WT and V5 hSIRT1 variants. Initial rates were measured by the continuous assay following incubation of the hSIRT1 variants with different concentrations of acetylated peptides, (**A**) H3K9Ac, (**B**) H3K56Ac, (**C**) H4K16Ac, (**D**) p53 containing K382Ac, (**E**) Primitive p53 containing 2 mutations in human p53 peptide and (**F**) RelA/p65 containing K310Ac. The initial rates were plotted against substrate concentration and fitted to the MM equation to derive the k_cat_ and K_M_ parameters reported in **Table S2**. Each experiment was performed in triplicate and representative experiment is shown. All peptide sequences are shown in **Table S3**.

**Fig. S9:**
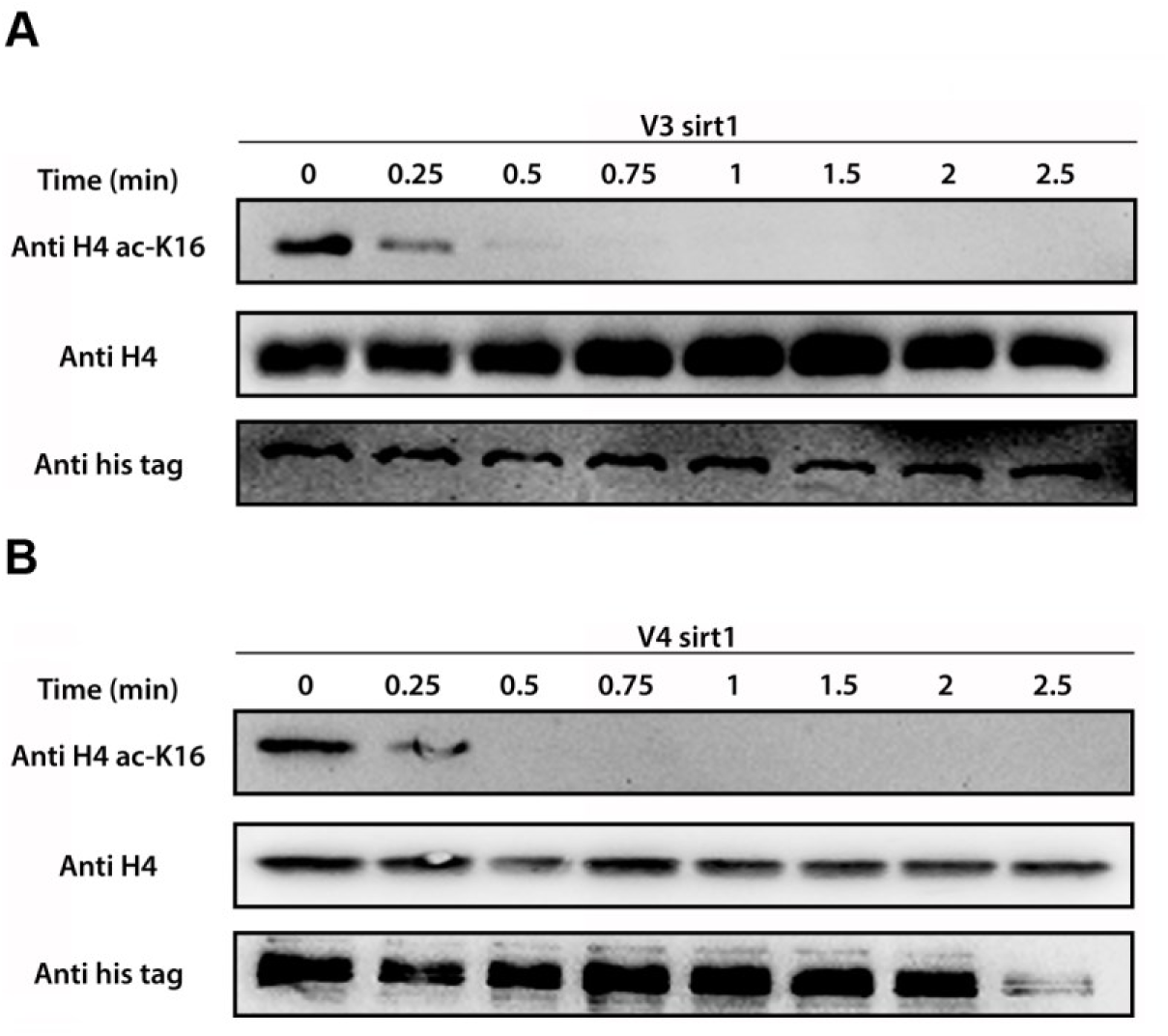
High V3 and V4 deacetylation activity toward native H4K16Ac. Western blot kinetic analysis of V3 (**A**) and V4 (**B**) activity toward H4K16Ac in the context of native histones. The H4K16Ac, H4 and hSIRT1 were detected using anti-H4K16Ac antibody, anti-H4 antibody and anti-6xHis antibody, respectively, as described in Methods section.

**Fig. S10:**
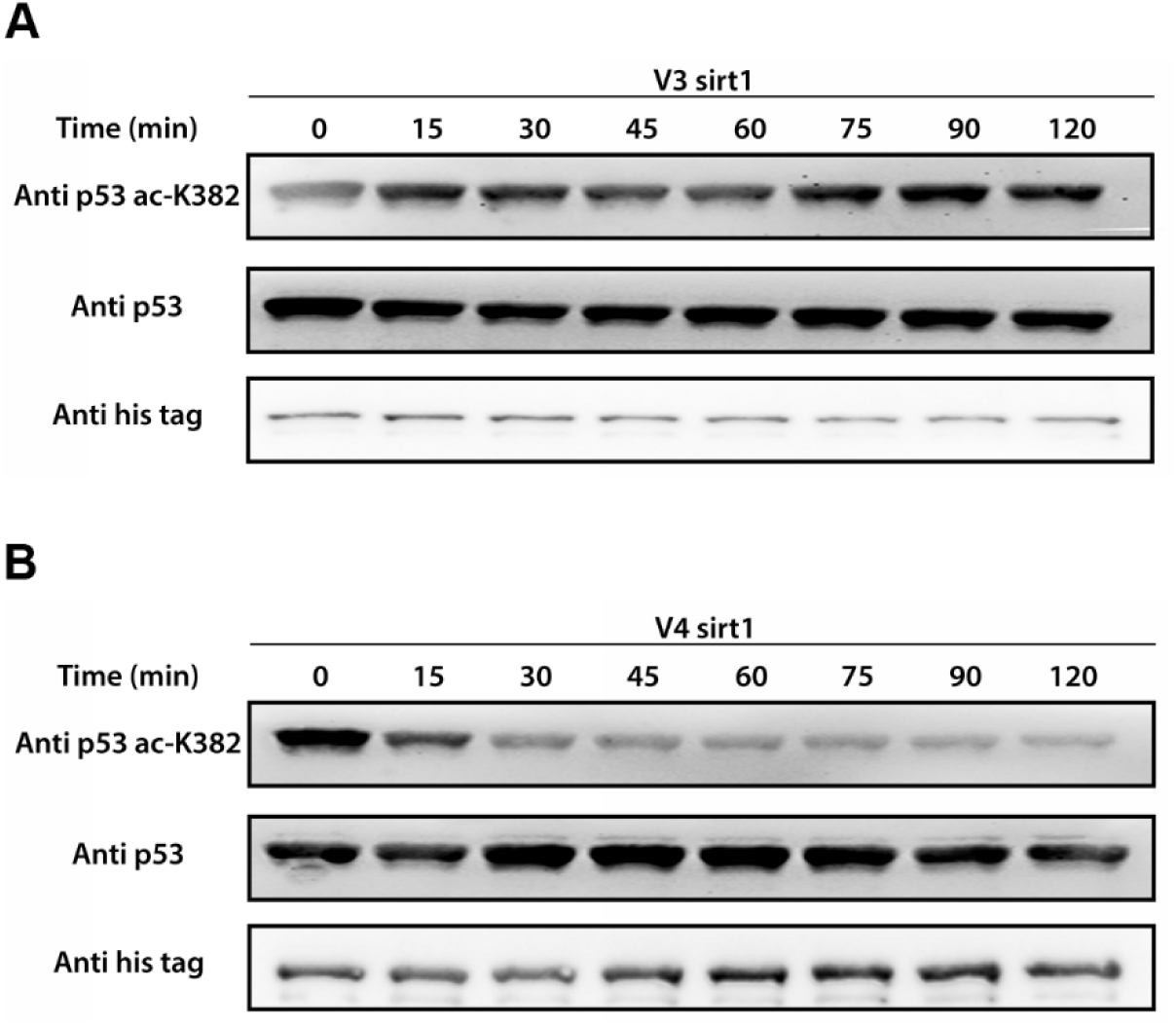
The activity of V3 and V4 with native acetylated p53. Representative western blot blots for the analysis of V3 (**A**) and V4 (**B**) activity with native p53 K382Ac. Detection of p53-K382Ac, p53 and hSIRT1 levels was performed using anti-K382Ac antibody, anti-p53 antibody and anti-6xHis antibody, respectively. Western blot kinetic analysis of V3 and V4 activity toward p53-K382Ac in the context of native p53 are shown in **Fig. 5**.

**Fig. S11:**
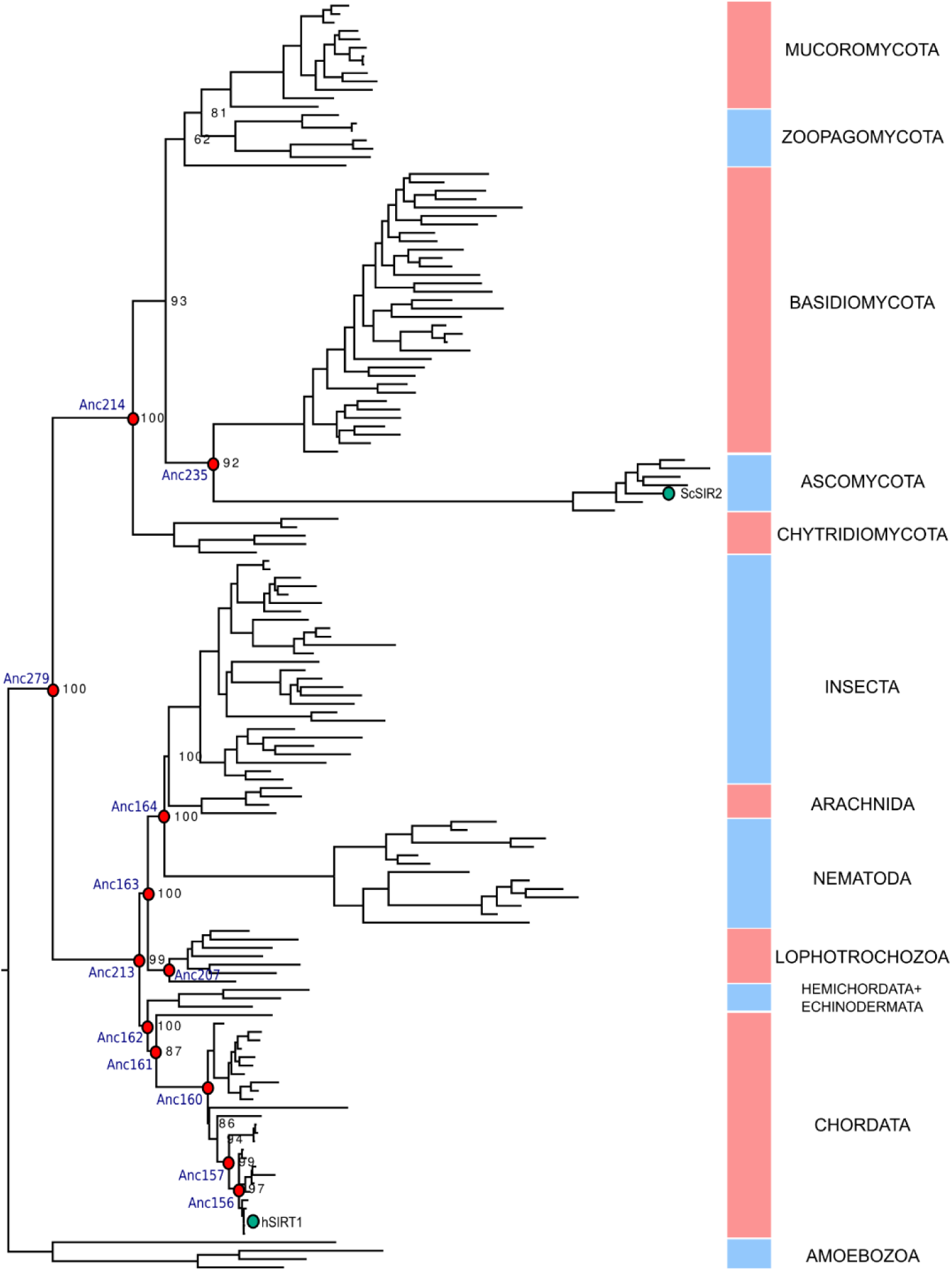
The constrained phylogeny based on a 151 sequence alignment of SIRT1 family inferred with IQ-TREE using the LG+I+G4 evolutionary model. Reconstructed nodes are labelled in red. The hSIRT1 and *Saccharomyces cerevisiae* Sir2 (ScSIR2) are labeled in green.

**Fig. S12:**
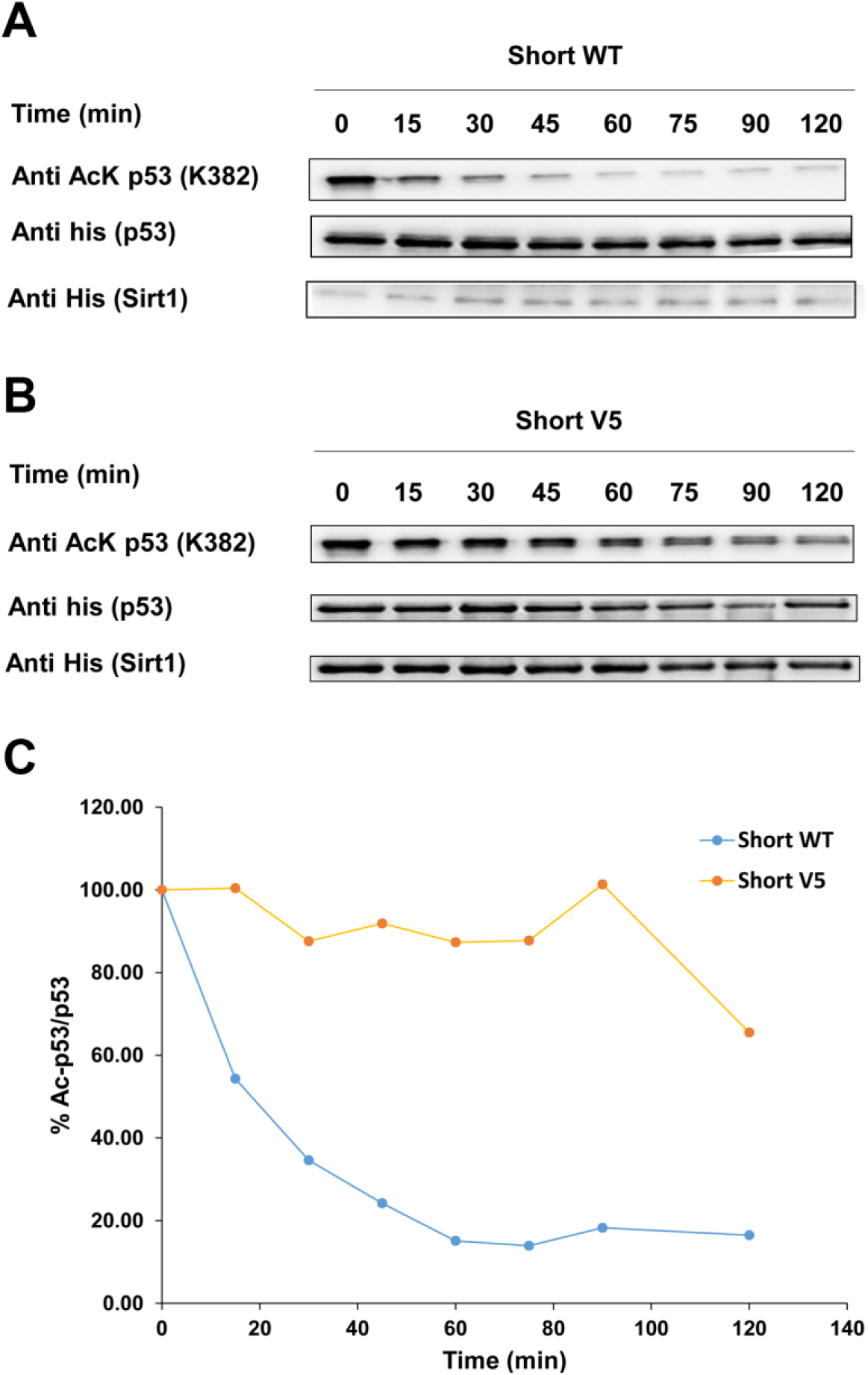
The activity of short truncated versions of WT and V5 with native acetylated p53. Western blot blots for the analysis of short WT (**A**) and short V5 variant (**B**) activity with native p53 K382Ac. Detection of p53-K382Ac, p53 and hSIRT1 levels was performed using anti-p53 K382Ac antibody and anti-6xHis antibody for p53 and Sirt1. (**C**) Western blot kinetic analysis of short WT and short V5 variant activity toward p53-K382Ac in the context of native p53. The time dependent decrease in K382Ac signal was normalized to time 0 and shown as the percentage of p53-K382Ac/p53 bands. Bands intensity were quantified by Image J.

**Fig. S13:**
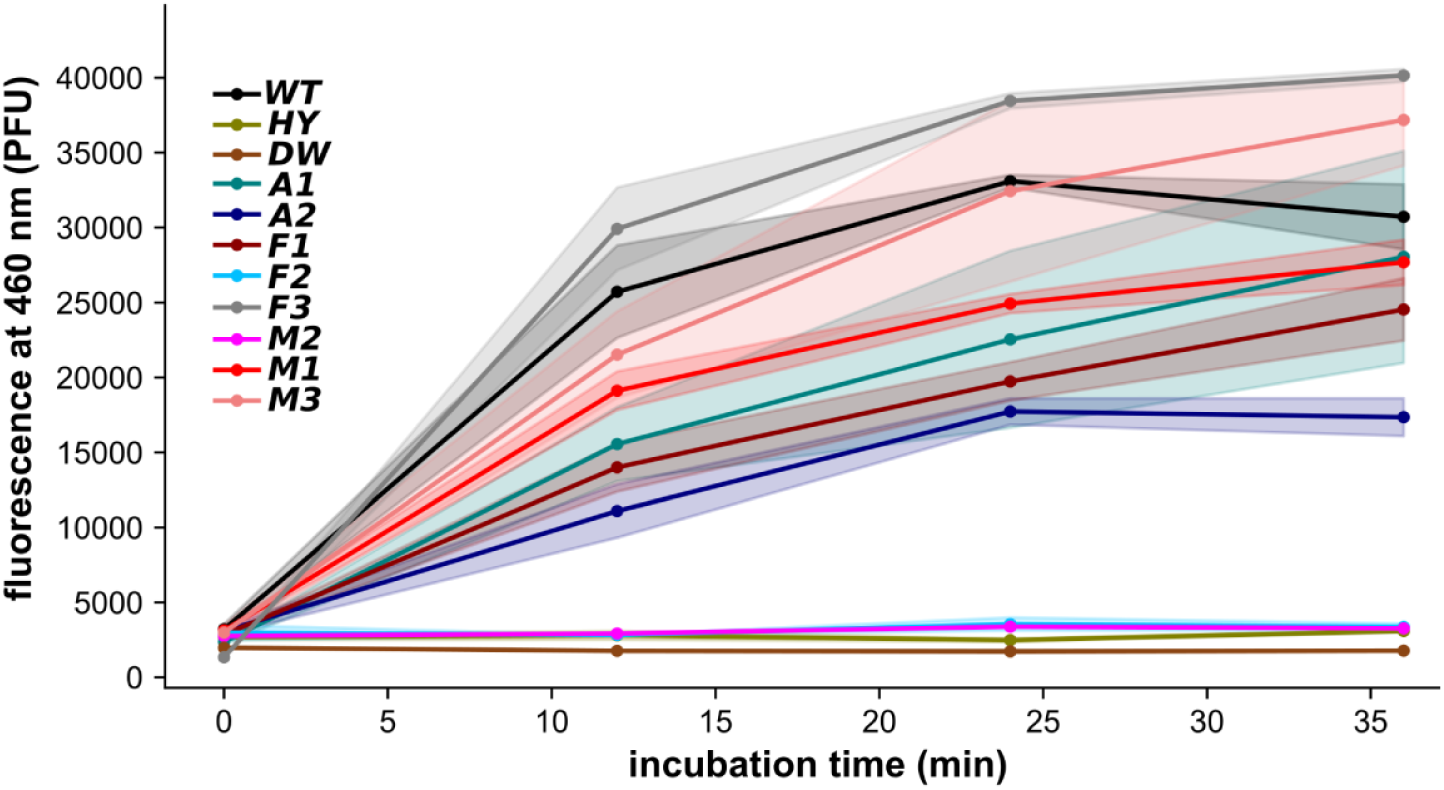
Fluor De lyse (FDL) analysis of hSIRT1 ASR mutants with protected acetyl-lysine-AMC substrate. Representative curves of fluorescent signal (PFU) at 460 nm for deacetylation of Boc-AcK-AMC. Shaded regions around curves show the standard error of the mean (SEM) for 2 technical replicates. WT served as the positive control; H363Y (HY; a mutation to the catalytically essential histidine at position 363) served as the the negative control, alongside DW (deionised water).

**Fig. S14:**
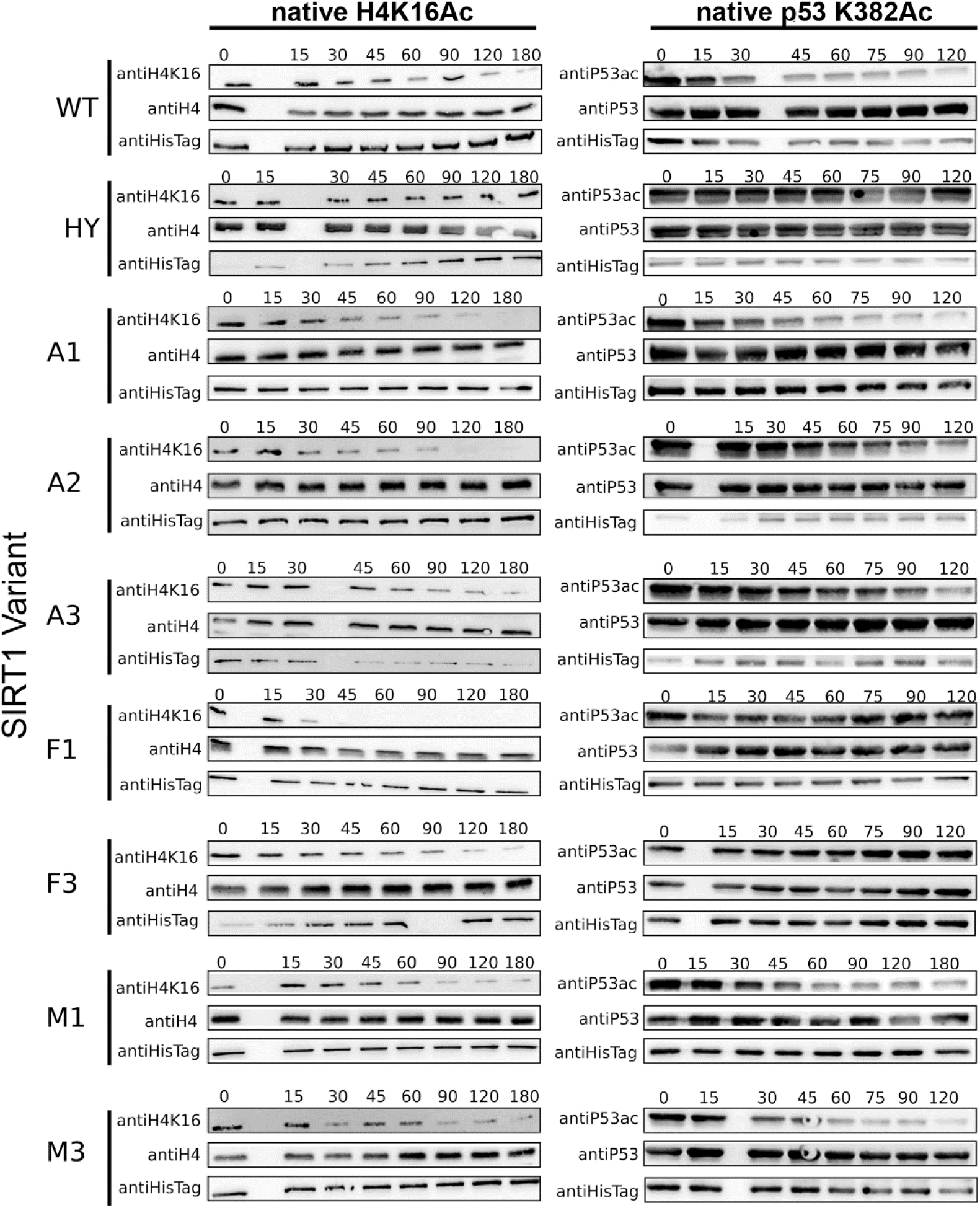
Representative western-blot analysis of SIRT1 ASR variants activities with native H4K16Ac and p53 K382Ac. Activity with histones (H4K16Ac) was measured up to 3 min (180 sec) of incubation while activity with p53 K382Ac was measured up to 120 min of incubation. All assays were performed as described in Methods and in experiments shown in Figs. 4-5.

**Table S1:** Documented hSIRT1 substrates and acetylation sites (analyzed in Fig. 1).

|  | **Substrate** | **Deacetylated lysine<sup>a</sup>** | **Reference (PMID)** |
| --- | --- | --- | --- |
| 1 | Histone H4 | K16 | 10693811 |
| 2 | Histone H3 | K9, K56 | 19879981, 19411844 |
| 3 | Tip60 | ND | 20100829 |
| 4 | PCAF | ND | 19188449 |
| 5 | ATG7 | ND | 25316028 |
| 6 | XPA/XPC | K67 | 20670893 |
| 7 | STK11(LKB11) | ND | 18687677 |
| 8 | MEF2 | K424 | 16166628 |
| 9 | Apex1 | K7 | 20699270 |
| 10 | Ku70 | K539/K542 | 15023334 |
| 11 | WRN | ND | 20428248 |
| 12 | MyoD | ND | 22771996 |
| 13 | p300/CBP | K1020 | 15632193 |
| 14 | HIF-1α | K647 | 20620956 |
| 15 | FOXO3 | ND | 21841822 |
| 16 | FOXO1 | ND | 15220471 |
| 17 | PPARγ | K268 | 22863012 |
| 18 | p53 | K382 | 11672523 |
| 19 | c-Myc | ND | 22190494 |
| 20 | RelA | K310 | 18419308 |
| 21 | IRS2 | ND | 18590691 |
| 22 | HIC1 | K314 | 17283066 |
| 23 | NBN(NBS1) | ND | 17612497 |
| 24 | CIITA | ND | 21890893 |
| 25 | PML | K487 | 22274616 |

<sup>a</sup>ND- non determined

**Table S2:** Kinetic parameters of WT and V3-5 with different acetylated peptides. Parameters were derived from fitting the kinetic measurements of the different hSIRT1 variants to the MM equation (for representatives kinetics for V5 see **Fig. S8**).

| **SIRT1** | **WT SIRT1** |  |  | **V3 SIRT1** |  |  | **V4 SIRT1** |  |  | **V5 SIRT1** |  |  |
| --- | --- | --- | --- | --- | --- | --- | --- | --- | --- | --- | --- | --- |
| **Parameters** | **$k_{cat}$**<br>**(sec<sup>-1</sup>)** | **$K_M$**<br>**(μM)** | **$k_{cat}/K_M$**<br>**(mM<sup>-1</sup>sec<sup>-1</sup>)** | **$k_{cat}$**<br>**(sec<sup>-1</sup>)** | **$K_M$**<br>**(μM)** | **$k_{cat}/K_M$**<br>**(mM<sup>-1</sup>sec<sup>-1</sup>)** | **$k_{cat}$**<br>**(sec<sup>-1</sup>)** | **$K_M$**<br>**(μM)** | **$k_{cat}/K_M$**<br>**(mM<sup>-1</sup>sec<sup>-1</sup>)** | **$k_{cat}$**<br>**(sec<sup>-1</sup>)** | **$K_M$**<br>**(μM)** | **$k_{cat}/K_M$**<br>**(mM<sup>-1</sup>sec<sup>-1</sup>)** |
| **Peptide** |  |  |  |  |  |  |  |  |  |  |  |  |
| **H4K16** | 0.2±<br>0.04 | 52.6±<br>18.5 | 5±2.3 | 0.3±<br>0.02 | 17.2±<br>4.4 | 20.3±5.8 | 0.3±0.02 | 19.8±<br>5.78 | 17.8±<br>5.04 | 0.2±0.01 | 8.8±3.7 | 26.8±5.2 |
| **H3K9** | 0.3±<br>0.07 | 101.6±<br>31.4 | 3.7±2.2 | 0.1±<br>0.04 | 17.6±<br>12.7 | 10.1±<br>3.3 | 0.1±0.05 | 36.5±22.<br>7 | 4.8±2.2 | 0.1±0.03 | 47±24.5 | 2.6±0 |
| **H3K56** | 0.1±<br>0.01 | 20.5±<br>7.6 | 7.5±2.3 | 0.1±<br>0.001 | 30.9±8.2 | 5.6±2.1 | 0.2±0.01 | 16.4±0 | 12.5±3.7 | 0.1±0.01 | 19.4±8.3 | 6.5±1.9 |
| **p53** | 0.4±<br>0.02 | 17.4±<br>3.8 | 25.8±<br>7.07 | 0.1±<br>0.01 | 10.9±3.5 | 15.7±3.5 | 0.2±0.03 | 26.2±<br>14.1 | 8.3±2.6 | 0.3±0.02 | 42.2±6.2 | 8.2±3.5 |
| **Primitive p53** | 0.3±<br>0.03 | 23.6±<br>7.4 | 15.5±<br>5.2 | 0.2±<br>0.04 | 10.4±9.4 | 22.6±<br>5.06 | 0.4±0.04 | 22.8±7.4 | 18.8±6.2 | 0.2±0.02 | 11.4±6.1 | 19.9±4.5 |
| **p65** | 0.1±<br>0.01 | 3.6±2.6 | 44.5±<br>4.6 | 0.2±<br>0.01 | 11±3.6 | 22.3±5.8 | 0.3±0.02 | 8.2±2.9 | 41.3±7.7 | 0.3±0.04 | 30.1±<br>12.8 | 9.9±3.7 |

**Table S3:**
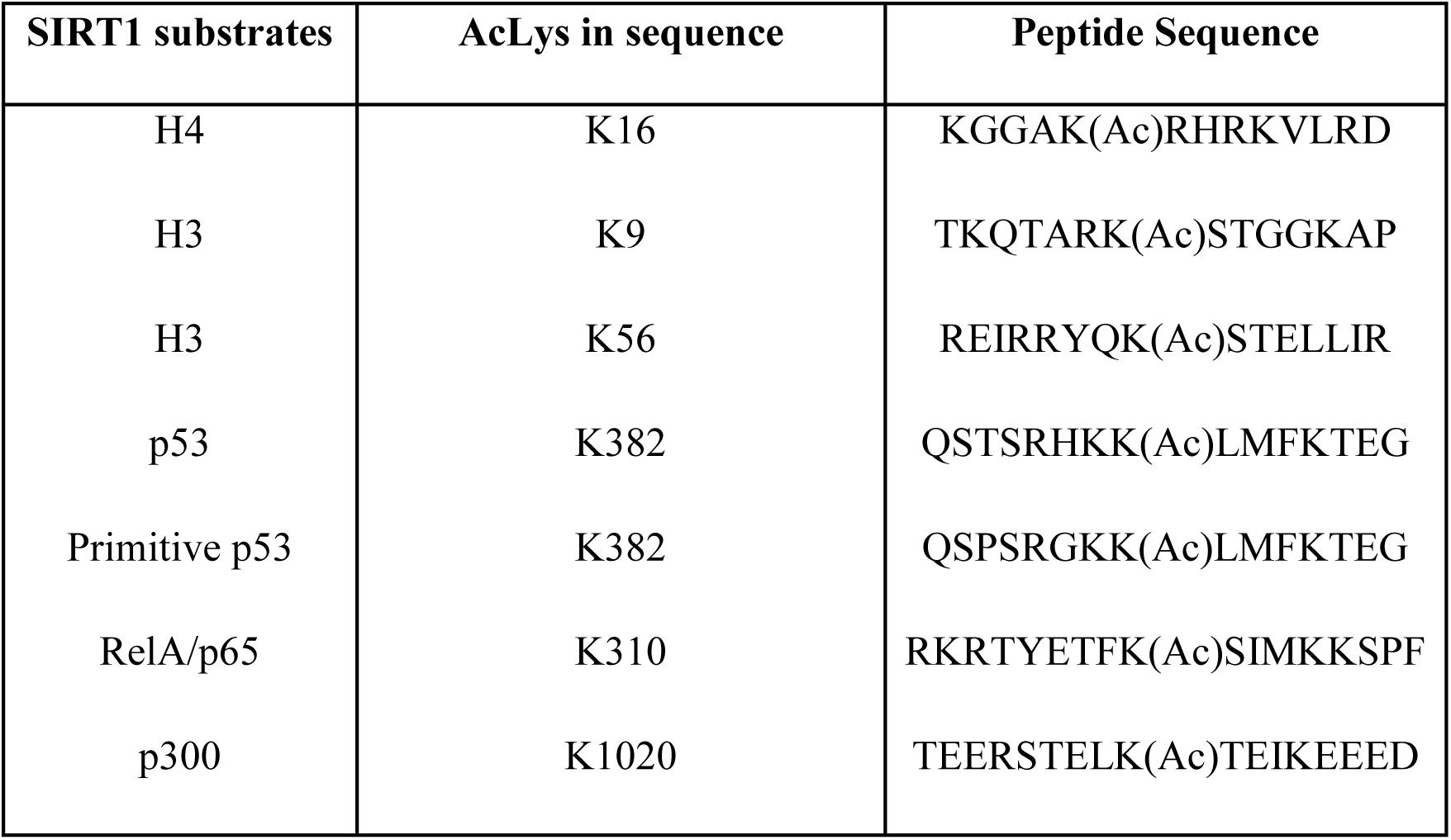
Peptide sequences of SIRT1 substrates used in this study.

| **SIRT1 substrates** | **AcLys in sequence** | **Peptide Sequence** |
| --- | --- | --- |
| H4 | K16 | KGGAK(Ac)RHRKVLRD |
| H3 | K9 | TKQTARK(Ac)STGGKAP |
| H3 | K56 | REIRRYQK(Ac)STELLIR |
| p53 | K382 | QSTSRHKK(Ac)LMFKTEG |
| Primitive p53 | K382 | QSPSRGKK(Ac)LMFKTEG |
| RelA/p65 | K310 | RKRTYETFK(Ac)SIMKKSPF |
| p300 | K1020 | TEERSTELK(Ac)TEIKEEED |

